# SARS-CoV-2 virus infection of *Peromyscus leucopus* demonstrates that infection tolerance is not limited to agents for which deermice are reservoirs

**DOI:** 10.64898/2026.03.13.711660

**Authors:** Ana Milovic, Johannes S. Gach, Ioulia Chatzistamou, Gema M. Olivarria, Thomas E. Lane, Donald N. Forthal, Alan G. Barbour

## Abstract

The North American deermouse *Peromyscus leucopus* is reservoir for several zoonotic agents, including bacterial, protozoan, and viral. It is remarkable for indiscernible or limited fitness consequences of these infections, a trait known as infection tolerance. But experimental infections have largely been of pathogens that *P. leucopus* naturally harbors. We asked whether infection tolerance extended to an agent, like SARS-CoV-2 virus, it had presumably not encountered before. Following protocols for experiments with mice and hamsters, we infected 8 female and 8 male *P. leucopus* of heterogeneous stock and compared responses of these animals on days 3 or 6 to those of 14 controls inoculated with virus-free medium. Serologic and virologic confirmation of infection was obtained for all exposed deermice. Moderate inflammation in lungs was histologically evident in infected animals, but no histological changes were noted in brains, even when viral RNA was present. Fourteen (88%) animals displayed no or only mild sickness; two had more severe illness. Genome-wide RNA-seq revealed an interferon-stimulated response on day 3 superceded mainly by a cell-mediated response by day 6. In brains transcription of the interferon-stimulated genes *Isg15* and *Mx2* positively correlated with viral RNA levels. The findings confirmed susceptibility of this species of *Peromyscus* to SARS-CoV-2 virus. For most infected outbred animals the immune response was swift and effective in controlling the pathogen and without evidence of excessive inflammation. Whatever is the basis for *P. leucopus’* trait of infection tolerance, it extended to at least one pathogen that for it would be novel.

**Importance:** *Peromyscus leucopus* is North American rodent that is reservoir for several agents of human disease, while exhibiting minimal illness, a phenotype termed infection tolerance. Whether this trait is pathogen-specific or represents a broader strategy has remained uncertain. By experimentally infecting *P. leucopus* with SARS-CoV-2 virus, which it is unlikely to have encountered, we investigated whether infection tolerance extends to a novel virus. Despite disseminated infection and lung pathology, most animals showed only mild or no disease. Expression analyses revealed early interferon-stimulated responses followed by cell-mediated responses with only limited production of inflammatory mediators interferon-gamma and nitric oxide synthase 2. Compared with results with a mouse model of infection, deermice displayed higher baseline expression of antiviral genes and quicker resolution of interferon responses. These findings suggest that infection tolerance is a strategy that limits immunopathology generally while resisting microbes, which has implications for understanding reservoir competence and host resilience.

## Introduction

Members of the cricetine rodent genus *Peromyscus*, commonly known as deermice (1), are widely distributed across North and Central America in varied wildland and peridomestic environments (2, 3). Deermice have been studied in evolutionary biology and ecological contexts and are experimental models for studies of infectious diseases, reproductive biology, behavior, and aging (4-7). One of these species, *P. leucopus*, the white-footed deermouse, is an important reservoir in North America for several agents of zoonoses, including Lyme disease, hard-tick relapsing fever (*Borrelia miyamotoi*), anaplasmosis, babesiosis, and deer tick or Powassan viral encephalitis (8, 9). Another species, *P. maniculatus*, is a reservoir for the Sin Nombre orthohantavirus (10, 11), the cause of hantavirus cardiopulmonary syndrome of humans, and the Modoc flavivirus (12). But this animal showed little sign of illness or disability from either virus (13, 14).

While several experimental infections of *P. leucopus* with the Lyme disease agent *Borreliella burgdorferi* are represented in the literature (15-22), there are fewer reports on experimental infections with other pathogens for which this species is a reservoir. Among the few studies were those of Mlera et al. of the Powassan encephalitis flavivirus, which can cause severe disease or death in human patients and experimentally-infected *Mus musculus* (23, 24). In these studies *P. leucopus* displayed little ill effect from systemic infection with the virus. The absence of evident sickness was also noted for *P. leucopus* experimentally infected with the Sin Nombre orthohantavirus, even when there were high virus titers in blood and organs (14). These findings added to the other examples of the phenomenon of infection tolerance in this species (8). By “infection tolerance” we mean immunological and physiological adaptations that minimize the harm from a pathogen’s presence (25, 26).

Aside from the aforementioned in vivo studies and an in vitro study of *P. leucopus* fibroblasts infected with viruses (27), there had been little attention on experimental infections of *P. leucopus* with a virus. What made a SARS-CoV-2 infection model feasible for this species and other members of the genus was an angiotensin-converting enzyme 2 (ACE2) protein that was suitable as a receptor for the virus (28). This was also case for the golden hamster (*Mesocricetus auratus*), another cricetine rodent and alternative animal model for pre-clinical SARS-CoV-2 studies (29). In this characteristic, deermice and hamsters were distinguished from *M. musculus*, whose ACE2 protein had substitutions that rendered it unsuitable as a receptor for this virus (30).

Confirmation of the susceptibility of *Peromyscus* species was provided in two contemporaneous reports of infections with SARS-CoV-2 virus of *P. maniculatus nebracensis* or *P. maniculatus rufinis* (31, 32). Following these was a report of infections of two other sub-species of *P. maniculatus* (*P. m. sonoriensis* and *P. m. bairdii*), *P. polionotus*, and the more distantly related *P. californicus* (33). The three articles reported isolation of virus from the lungs and oral swabs, lung pathology, and the development of neutralizing antibodies in inoculated animals. Of the 3 species (including all 4 sub-species of *P. maniculatus*) studied, only *P. californicus* displayed signs of moderate to severe sickness in some of the inoculated animals (33). There had been serological evidence of betacoronaviruses infections in wild populations of *Peromyscus* (34), but these viruses are divergent from the SARS-CoV-2 virus. With an assay specific for SARS-CoV-2 virus, another investigation confirmed lack of evidence of SARS-CoV-2 infections among *Peromyscus* populations in the eastern United States (35).

The prior studies of the *Peromyscus* species infected with SARS-CoV-2 virus justifiably focused on the potential for these commonly encountered animals to be sources of transmission of the virus to humans and other mammals in North America. Two of the studies also included analyses of expression of selected genes, but these were limited to the lungs and in the number of genes examined (31, 32). The present study aimed, first, to extend this infection model to *P. leucopus*, the only species for which there was available a high-coverage, chromosome-scale genome assembly with annotation based on transcriptomes of several organs and tissues, not just predictions of transcripts (36). The closed colony population from which the animals would be drawn has genetic diversity that is close to that of natural populations of the species and in this characteristic comparable to most human populations (37). If this first aim succeeded, we would then expand the sample sizes, match virus-exposed animals with controls inoculated with virus-free medium, broaden the transcriptomic analysis to genome-wide in scope, and include the brain as well as lungs. These would complement virologic, serologic, and histopathologic characterizations of the experimental infections.

The study would also apply a test to a hypothesis about infection tolerance in *Peromyscus*. If *P. leucopus* tolerates the infections by pathogens for which it is natural host, this conceivably could be attributable to cumulative more-or-less singular adaptations over an extended time, such that tolerance of, for example, *B. burgdorferi* infection was specific to that bacterium. Whatever that particular accommodation entailed, it would not necessarily account for *P. leucopus*’ adaptation to infection with the protozoan *Babesia microti* or with the flavivirus of Powassan encephalitis. If this was the history of its evolution, then infection with SARS-CoV-2 virus, a novel pathogen for *P. leucopus*, might be more severe in that species than experience to date would predict. On the other hand, if we are looking at a trait, likely multigenic in basis, that confers broader capacity for infection tolerance, then this could be as applicable for newly-threatening pathogens as ones the deermice have been living with for ages.

## Results

### Experimental infections

Adult *P. leucopus* of both sexes were inoculated intranasally on day 0 with either SARS-CoV-2 virus in culture medium or, as a control, medium alone in the same volume (Figure 1). Virus-inoculated animals in a BSL-3 facility were housed separately from controls. There were two replicates (“experiment 1” and “experiment 2”) using different stocks of the same SARS-CoV-2 USA-WA1 strain but otherwise under the same conditions (Table 1). The mean and median ages for the subject animals over both experiments were 582 days and 578 days, respectively, with range of 219-998 days. As expected, males (mean of 22 g) were overall ∼15% larger in body mass than females (mean of 19 g) at entry (*p* = 0.01), but there was no apparent association between age and body mass across sexes (*R*^*2*^ = 0.04). Given the maximum life span of 8 years for the species (38), even those between 2-3 years old would not be considered aged animals. Animals were monitored daily for signs of distress, weight loss, and sickness behavior with assignment of a sickness score of 0, 1, 2, or 3. On day 3 or 6 the animals were euthanized, and specimens of blood, lung, and brain were collected for histopathology, RNA extraction, virus detection, and antibody titers.

**Figure 1.**
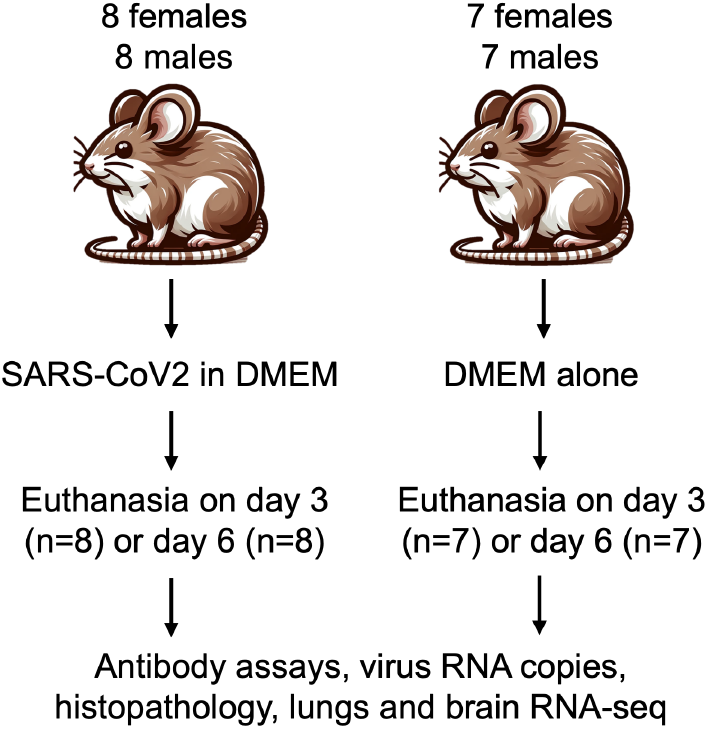
Experimental design and course for intranasal inoculation of female and male adult *Peromyscus leucopus* with SARS-CoV-2 virus (2×10^4^ particles) or DMEM culture medium alone and the collection of specimens and euthanasia on either day 3 or day 6 after inoculation.

**Table 1.** Line listing of animals inoculated with SARS-CoV-2 or medium alone by their characteristics, viral burden, and antibody responses.

|  |  |  |  |  |  |  |  |  |  |  |  |  |  |  |  |
| --- | --- | --- | --- | --- | --- | --- | --- | --- | --- | --- | --- | --- | --- | --- | --- |
| Exp* | Animal | Sex* | Age (d) | Body mass (g) at day 0 | Inoculum | Euthanasia day | Sickness score | Lung path. score | Log<sub>10</sub> lung viral RNA copies | 1/anti-S titer | 1/anti-N titer | 1/neutralization titer | Log<sub>10</sub> brain viral RNA copies | Log<sub>10</sub> virus reads brain RNA-seq | Log<sub>10</sub> feces viral RNA copies |
| 1 | 25160 | F | 501 | 22 | medium | 3 | 0 | 0 | 0.0 | <50 | 381 | 1 | 2.6 | 0.3 | <2.5 |
| 1 | 25169 | M | 483 | 19 | medium | 3 | 0 | 0 | 1.3 | <50 | 100 | 1 | 2.1 | 0.6 | <2.5 |
| 1 | 25124 | M | 578 | 20 | medium | 3 | 0 | 0 | 2.2 | <50 | 113 | 1 | 1.1 | 0.8 | <2.5 |
| 1 | 25167 | F | 486 | 22 | medium | 6 | 0 | 0 | 2.2 | 598 | 120 | 1 | 2.1 | 0.7 | <2.5 |
| 1 | 25018 | F | 627 | 19 | medium | 6 | 0 | 0 | 0.0 | <50 | <50 | 1 | 1.8 | 0.6 | <2.5 |
| 1 | 25159 | M | 499 | 20 | medium | 6 | 0 | 0 | 0.0 | <50 | 458 | 1 | 2.2 | 0.6 | <2.5 |
| 1 | 25013 | F | 662 | 18 | virus | 3 | 0 | 1 | 6.5 | <50 | 736 | 1 | 4.3 | 1.6 | 6.2 |
| 1 | 25025 | F | 761 | 19 | virus | 3 | 0 | 1 | 7.1 | <50 | 156 | 1 | 3.0 | 0.3 | 2.7 |
| 1 | 25117 | M | 583 | 22 | virus | 3 | 0 | 2 | 6.8 | <50 | 205 | 1 | 4.1 | 1.7 | 5.6 |
| 1 | 25130 | M | 578 | 19 | virus | 3 | 0 | 2 | 7.7 | <50 | 300 | 1 | 4.8 | 2.5 | 3.4 |
| 1 | 25128 | F | 581 | 19 | virus | 6 | 0 | 1 | 4.9 | 38632 | 643 | 459 | 4.6 | 2.3 | 3.1 |
| 1 | 25158 | F | 504 | 17 | virus | 6 | 0 | 1 | 3.1 | 30203 | 161 | 186 | 3.6 | 1.9 | <2.5 |
| 1 | 25157 | M | 504 | 29 | virus | 6 | 0 | 1 | 5.1 | 35470 | 730 | 108 | 2.4 | 0.7 | 4.1 |
| 1 | 25161 | M | 509 | 26 | virus | 6 | 0 | 1 | 5.6 | 56519 | 1914 | 334 | 2.0 | 0.5 | <2.5 |
| 2 | 25225 | F | 287 | 16 | medium | 3 | 0 | 0 | 2.1 | <50 | 182 | 1 | 3.5 | 0.0 | . |
| 2 | 25226 | F | 223 | 20 | medium | 3 | 0 | 0 | 0.0 | 96 | 474 | 1 | 2.8 | 0.8 | . |
| 2 | 25281 | M | 644 | 21 | medium | 3 | 0 | 0 | 0.6 | <50 | 160 | 1 | 1.3 | 0.0 | . |
| 2 | 25152 | M | 949 | 24 | medium | 3 | 0 | 0 | 0.0 | 79 | 253 | 1 | 2.4 | 0.6 | . |
| 2 | 25219 | F | 317 | 17 | medium | 6 | 0 | 0 | 0.6 | <50 | 376 | 1 | 3.0 | 0.7 | . |
| 2 | 25313 | F | 373 | 16 | medium | 6 | 0 | 0 | 1.3 | <50 | 477 | 1 | 3.0 | 1.0 | . |
| 2 | 25287 | M | 675 | 21 | medium | 6 | 0 | 0 | 0.0 | <50 | 268 | 1 | 1.5 | 1.0 | . |
| 2 | 25280 | M | 647 | 17 | medium | 6 | 0 | 0 | 0.0 | <50 | 167 | 1 | 1.2 | 0.7 | . |
| 2 | 25298 | F | 626 | 17 | virus | 3 | 1 | 1 | 2.7 | 121 | 2133 | 1 | 6.7 | 2.8 | . |
| 2 | 25304 | F | 469 | 19 | virus | 3 | 1 | 2 | 3.8 | 220 | 2660 | 1 | 7.1 | 3.7 | . |
| 2 | 25255 | M | 932 | 21 | virus | 3 | 3 | 1 | 6.8 | 126 | 577 | 1 | 7.2 | 3.9 | . |
| 2 | 25165 | M | 919 | 19 | virus | 3 | 3 | 2 | 5.4 | 460 | 2364 | 1 | 6.4 | 2.7 | . |
| 2 | 25306 | F | 472 | 16 | virus | 6 | 0 | 2 | 5.6 | 27066 | 2515 | 136 | 6.6 | 3.1 | . |
| 2 | 25228 | F | 219 | 24 | virus | 6 | 0 | 1 | 3.4 | 117372 | 2693 | 607 | 7.7 | 4.2 | . |
| 2 | 25136 | M | 952 | 23 | virus | 6 | 1 | 1 | 4.2 | 11500 | 2844 | 1 | 7.8 | 4.7 | . |
| 2 | 25163 | M | 922 | 22 | virus | 6 | 1 | 1 | 4.2 | 62542 | 761 | 164 | 5.8 | 2.5 | . |

\* F, female; M, male

None of the 14 animals inoculated with medium alone in either experiment 1 or 2 exhibited signs of distress or sickness behavior over the 3- or 6-day periods of observation. We similarly recorded absence of distress or sickness for the 8 animals that received virus in experiment 1. In contrast, 6 of the 8 animals receiving virus in experiment 2 were identified as overtly sick (Fisher exact *p* = 0.007): four with mild illness and score of 1 and two, animals 25165 and 25255, with scores of 3 (Table 1). Manifestations of mild illness were piloerection and periorbital edema. Those with severe illness in addition displayed lethargy and tachypnea by day 2 and were euthanized based on humane endpoints on day 3. Four of the 6 ill animals, including both severely ill animals, were older than 900 days. The two deermice that did not exhibit sickness behavior in experiment 2 were female and 219 or 472 days of age. In experiment 2 change of weight over 3 or 6 days was monitored for 8 control and 8 virus-infected animals; the mean (95% confidence interval [CI]) body mass change +0.25 (-0.08 to +0.58) g for controls and +0.18 (-0.87 to +1.22) g for infected deermice (*p* = 0.89).

Cultivation of lung tissue from day 3 by focus forming assay was carried out for specimens of two experiment 1 animals, 25025 and 25117, and yielded 30 PFU and 4 PFU per mg of lung tissue, respectively, thereby confirming presence of infectious virus in the lung. Viral RNA copies, as assessed by RT-qPCR, varied over a 10^4^-fold range in both lung and brain (Table 1 and Figure 2). By day 6 viral copy numbers in the lung were overall ∼100-fold lower than on day 3. There was a greater range of viral RNA copies values for the brain tissue than for lung on day 6 and, unlike the lung, little difference in the ranges of copy numbers between day 3 and day 6 specimens. The four highest copy numbers of viral RNA for both day 3 and day 6 brain were from experiment 2. The coefficient of determination (*R*^*2*^) for RT-qPCR viral RNA copies in lungs and brains was 0.30 (*p* <0.001). For experiment 1 RNA was also extracted from fecal pellets collected from individual animals at the time of euthanasia; virus was detected at limit of sensitivity of the RT-qPCR assay in 6 of the 8 virus-inoculated animals and none of the fecal samples from medium-inoculated controls (Table 1). Female and male deermice were not distinguishable in their viral burdens in lung, brain, or feces. There was no apparent association between age of animal at entry and either lung (*R*^*2*^ = 0.05) or brain (*R*^*2*^ = 0.04) viral RNA copies.

**Figure 2.**
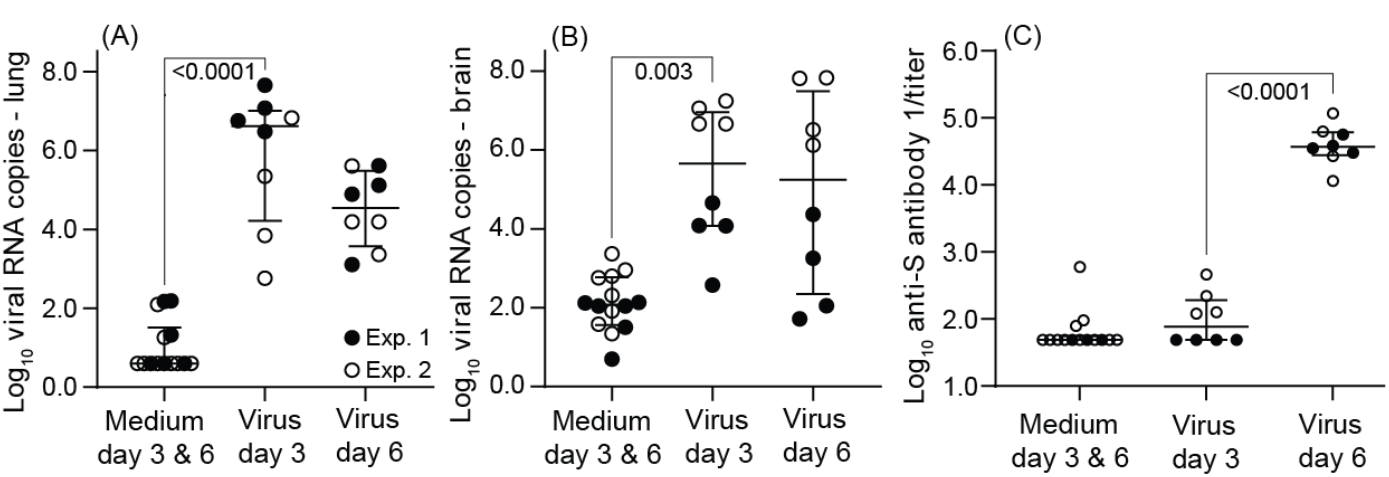
Box-whisker plots of log-transformed values for viral RNA burden in lung (panel A) and brain (panel B) and anti-spike protein (S) antibody titers (panel C) in SARS-CoV-2-infected and control *P. leucopus* on either day 3 or day 6 in experiments (Exp) 1 and 2. Anti-spike (S) antibody titers were measured by ELISA and calculated by endpoint titers. Box-whisker plots show mean and 95% confidence intervals. One-way analysis of variance (ANOVA) test was used to analyze the significant differences in viral load and titers between different groups. Data for the graphs are from Table 1.

All virus-inoculated animals had detectable anti-SARS-CoV-2 Spike (S) antibodies by day 6, and low levels were also discernible in all day 3 blood specimens from experiment 2 animals (Table 1 and Figure 2). Anti-SARS-CoV-2 Nucleocapsid (N) antibody titers were generally lower than anti-S antibody titers across both experiments, and they were not significantly different from control values for either day 3 or day 6 for experiment 1 animals. In contrast, for experiment 2 plasma samples the mean (95% CI) anti-N titers were generally higher on both day 3 [1934 (1004-2863); *p* = 0.0004] and day 6 [2203 (1232-3174); *p* = 0.0002] than for day 3 and 6 controls [295 (201-388)]. Neutralizing antibody titers above background were only observed in blood samples obtained on day 6 (Table 1).

In summary, both experiments demonstrated by virologic and serologic criteria that this population of *P. leucopus* can be infected with a human isolate of the SARS-CoV-2 virus and that for only a minority of animals was the disease more than mild in severity. However, it was also evident that the different stocks of the USA-WA1 strain for the two experiments differed in the infections they caused. There was an earlier antibody response, higher virus burden in the brain, and higher proportion of animals observably sick in experiment 2.

### Histopathology of lung and brain

Specimens from deermice infected with the virus showed changes consistent with acute lung injury on both day 3 and day 6 by hematoxylin and eosin (H & E) staining (Table 1 and Figure 3). Signs of acute inflammation were marginally more pronounced on day 3 of infection; 4 of the 5 specimens with pathology scores of 2 were from day 3. Notable findings were mild focal bronchial submucosal inflammatory infiltrates, alveolar septal thickening and neutrophilic infiltrates, alveolar exudates, and hyaline membranes lining the alveoli (Figure 3 panels E-G). The lungs of the two infected animals with a sickness score of 3 (Table 1) also showed evidence of alveolar hemorrhage, which was most prominent in animal 25165 (Figure 3 panel E). Several alveoli were filled with red blood cells, and there were congested capillaries, suggestive of early microthrombotic changes. The presence of virions in the lung tissue in an animal on day 3 of infection was demonstrated by in situ hybridization with a specific probe for virus (Figure S1).

**Figure 3.**
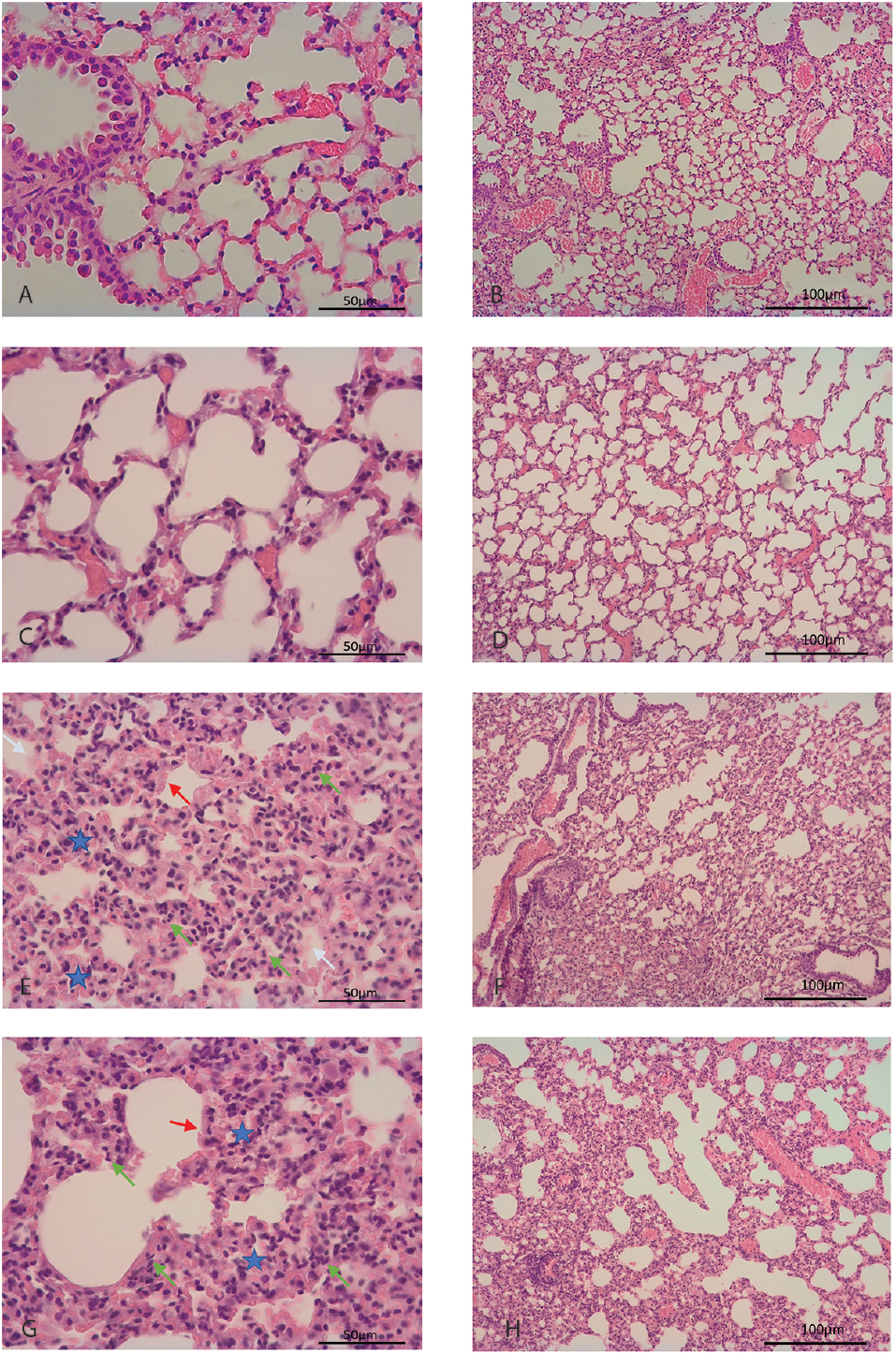
Photomicrographs of hematoxylin and eosin-stained lung sections of *P. leucopus* with and without infection by SARS-CoV-2 virus. These are displayed at low magnification (panels B, D, F, and H with 100 μm size bars) or high magnification (panels A, C, E, G with 50 μm size bars). Panels A and B are of an animal without any treatment. Panels C and D are of lung of an animal 3 days inhalation of medium alone. Panels E and F are of lung of a virus-infected animal at day 3. Panels G and H are of lungs of virus-infected animal at day 6. In panels E and G arrowheads indicate neutrophil infiltration (green), hyaline membranes (red), and alveolar exudate (white). A blue star denotes thickening of a septum.

Lung tissues from control animals, which had inhaled medium without virus, exhibited on days 3 or 6 mild intraseptal inflammation, characterized as a focal foreign body reaction, which was not observed in the lung from an untreated animal. Alveolar exudates, hemorrhage, or hyaline membranes were not observed in specimens that received medium alone. Use of these particular controls for RNA-seq studies allowed discernment of the specific contributions of the inhalation of virus over tissue reactions to the medium’s constituents alone.

H & E-stained sections of the brain, including the meninges, were assessed for inflammation, and features such as tissue edema, lymphocyte infiltration, perivascular hemorrhage, and intravascular thrombosis were evaluated. None of these features were observed in analyzed sections of any of the infected animals on either day 3 or day 6.

### RNA-seq of lung tissues

In the absence for *P. leucopus* of many of the reagents needed for flow cytometry or assays for specific proteins, bulk genome-wide, RNA-seq serves as a source for insights for this non-model organism (39, 40). For the lung tissues in the present study the mean (95% CI) number of 100 nt paired-end (PE100) reads of reverse-transcribed mRNAs were 6.8 (5.9-7.6) × 10^7^ for 30 samples. The reads were aligned to a reference set comprising 22,598 non-redundant coding sequences (CDS) from the reference *P. leucopus* genome GCF_004664715.2 (accession), which was further manually annotated (41).

Differential gene expression analysis comparing infected to control animals from both experiments revealed many more up-regulated than down-regulated genes in infected animals on each of the days (Figure 4, Table 2, and Table S1). By the criterion of fold change ≥2.0 up or down and a false-discovery rate *p* value <0.05, there were 135 up-regulated differentially-expressed genes (DEG) on day 3 and 31 on day 6. At 3 days there were also 7 genes, including the sodium channel *Scn7a* (NAX), lower in transcription than in controls by 2-3 fold, but there were no down-regulated DEGs on day 6 by those cut-offs.

**Figure 4.**
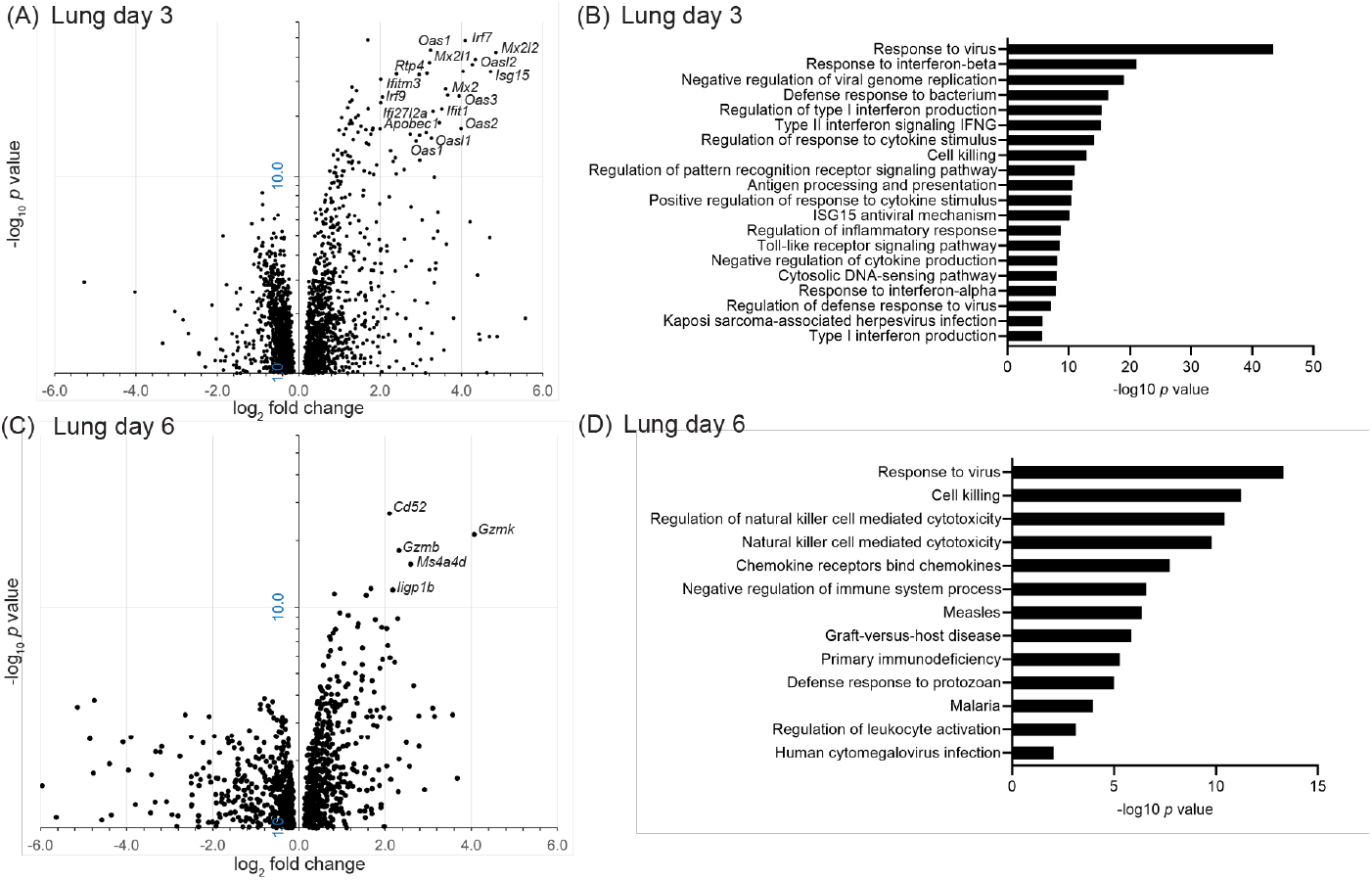
Volcano plots (panels A and B) and Gene Ontology (GO) term clusters (panels C and D) of bulk, genome-wide RNA-seq analysis lungs of *Peromyscus leucopus* on day 3 or day 6 after intranasal inoculation with SARS-CoV-2 virus or medium alone. The data for the volcano plots are provided in Table S1. In the volcano plot the *y*-axes are log_10_ scale, and the majorities of up-regulated genes with log_2_-fold change of ≥2.0 and false discovery rate *p*-values of <10^-10^ are indicated. The GO term analyses are for up-regulated pathways and functions; the identification numbers for each of the listed terms are given in Table S2.

**Table 2.** Up-regulated differentially expressed genes in lung by group (infected or control and day of infection)

|  |  |  |  |  |  |  |
| --- | --- | --- | --- | --- | --- | --- |
| Group 1 | Group 2 | Gene | Group 1 mean (95CI)* | Group 2 mean (95CI) | FC 2 vs 1 | FDR p value* |
| Control | Day 3 | *Apobec1* | 0.6 (0.4–0.8) | 6.7 (5.6–7.8) | 9.6 | 1E-13 |
|  |  | *Apol6* | 0.3 (0.2–0.4) | 1.1 (0.6–1.9) | 6.8 | 2E-02 |
|  |  | *Ccl3* | 0.6 (0.5–0.9) | 1.3 (0.5–3.6) | 9.3 | 4E-02 |
|  |  | *Cxcl11* | 0.5 (0.3–0.8) | 2.9 (1.8–4.7) | 8.2 | 2E-04 |
|  |  | *Gbp11* | 5.2 (4.5–6.0) | 39 (24–64) | 9.9 | 6E-19 |
|  |  | *Gm4841* | 4.7 (3.7–6.1) | 31 (20–50) | 7.8 | 4E-14 |
|  |  | *Gstm3* | 1.2 (1.0–1.6) | 2.8 (0.9–9.0) | 18.5 | 2E-04 |
|  |  | *Gzmb* | 0.6 (0.4–0.8) | 3.8 (2.3–6.3) | 6.8 | 2E-05 |
|  |  | *Ifi2712a* | 252 (162–391) | 2218 (1588–3098) | 7.9 | 3E-10 |
|  |  | *Ifit1* | 0.1 (0.1–0.2) | 1.0 (0.7–1.6) | 12.4 | 4E-03 |
|  |  | *ligp1b* | 2.7 (2.1–3.6) | 57 (40–80) | 20.3 | 8E-36 |
|  |  | *Irf7* | 17 (14–19) | 242 (162–363) | 17.1 | 6E-45 |
|  |  | *Isg15* | 21 (15–29) | 547 (362–826) | 26.3 | 7E-31 |
|  |  | *Itih3* | 3.7 (3.0–4.7) | 5.3 (1.7–16) | 10.7 | 8E-04 |
|  |  | *Mx2* | 1.4 (1.0–2.0) | 42 (29–61) | 28.9 | 4E-39 |
|  |  | *Oas1* | 0.3 (0.2–0.6) | 3.8 (2.7–5.4) | 10.0 | 3E-08 |
|  |  | *Oas2* | 0.5 (0.3–0.8) | 7.6 (6.1–9.4) | 11.0 | 2E-16 |
|  |  | *Oas3* | 0.7 (0.5–1.1) | 13 (9.4–18) | 15.4 | 8E-23 |
|  |  | *Oasl1* | 1.0 (0.7–1.4) | 10 (7.1–15) | 8.8 | 1E-14 |
|  |  | *Oasl2* | 1.3 (0.9–1.8) | 28 (21–389) | 19.2 | 1E-33 |
|  |  | *Rsad2* | 0.8 (0.6–1.0) | 11 (6.4–20) | 15.8 | 3E-15 |
|  |  | *Tgtp2* | 0.3 (0.2–0.4) | 2.0 (1.2–3.2) | 10.7 | 5E-05 |
|  |  | *Tnnt3* | 0.5 (0.3–0.9) | 1.2 (0.3–4.7) | 25.9 | 2E-03 |
|  |  | *Usp18* | 5.8 (5.1–6.6) | 49 (35–68) | 9.3 | 2E-34 |
| Control | Day 6 | *Ccl5* | 13 (10–18) | 51 (31–83) | 4.1 | 1E-05 |
|  |  | *Cd52* | 10 (8.0–12) | 44 (37–53) | 4.3 | 5E-23 |
|  |  | *Cd8a* | 6.4 (4.9–8.4) | 16 (13–19) | 2.1 | 2E-02 |
|  |  | *Ctsw* | 2.5 (2.0–3.1) | 7.3 (5.3–10) | 2.8 | 3E-04 |
|  |  | *Cxcl10* | 3.6 (2.8–4.6) | 12 (9.0–17) | 3.4 | 3E-06 |
|  |  | *Cxcl9* | 2.0 (1.6–2.6) | 7.4 (4.6–12) | 3.8 | 1E-03 |
|  |  | *Cxcr3* | 0.9 (0.6–1.2) | 5.5 (4.2–7.1) | 4.9 | 2E-06 |
|  |  | *Gbp11* | 5.2 (4.5–6.0) | 15 (12–20) | 2.9 | 1E-08 |
|  |  | *Gm4841* | 4.7 (3.7–6.1) | 12 (8–17) | 2.5 | 6E-03 |
|  |  | *Gzma* | 11 (8.4–14) | 38 (31–45) | 3.2 | 3E-09 |
|  |  | *Gzmb-2* | 2.1 (1.6–2.7) | 12 (10–15) | 5.0 | 7E-15 |
|  |  | *Gzmk* | 0.4 (0.3–0.7) | 12 (8.2–17) | 16.9 | 6E-18 |
|  |  | *ligp1b* | 2.7 (2.1–3.6) | 14 (9.9–20) | 4.5 | 4E-09 |
|  |  | *Klrk1* | 1.7 (1.4–2.0) | 5.3 (4.4–6.5) | 2.8 | 1E-02 |
|  |  | *IGHV5-4* | 6.3 (4.3–9.4) | 21 (13–34) | 3.2 | 1E-02 |
|  |  | *Trbv23* | 0.6 (0.3–0.9) | 3.3 (2.4–4.5) | 4.7 | 2E-03 |
|  |  | *IGHV1-55* | 11 (7.0–18) | 63 (39–102) | 4.3 | 1E-03 |
|  |  | *Ms4a4d* | 1.3 (0.9–1.7) | 9.0 (6.9–12) | 6.0 | 1E-12 |
|  |  | *Nkg7* | 7.4 (6.1–8.9) | 20 (16–26) | 2.6 | 6E-06 |
|  |  | *Oas1* | 0.3 (0.2–0.6) | 2.2 (1.0–4.6) | 6.3 | 2E-02 |
|  |  | *Pclaf* | 7.3 (5.6–9.5) | 22 (19–25) | 2.6 | 1E-05 |
|  |  | *Prf1* | 2.7 (2.1–3.6) | 7.6 (5.3–11) | 2.8 | 8E-03 |
|  |  | *Sh2d1a* | 2.0 (1.5–2.7) | 6.1 (4.4–8.6) | 2.7 | 3E-02 |
|  |  | *Sh2d2a* | 1.4 (1.1–1.9) | 6.2 (4.6–8.2) | 3.8 | 1E-05 |
|  |  | *Tap1* | 19 (17–22) | 42 (32–56) | 2.2 | 1E-06 |
| Day 3 | Day 6 | *Ca4* | 48 (41–56) | 93 (75–115) | 1.8 | 1E-02 |
|  |  | *Cd247* | 5.9 (5.0–6.8) | 13 (10–16) | 2.1 | 1E-02 |
|  |  | *Cd3e* | 8.8 (6.7–12) | 22 (17–27) | 2.2 | 2E-03 |
|  |  | *Cd3g* | 19 (16–24) | 43 (34–53) | 2.0 | 7E-03 |
|  |  | *Cd52* | 13 (9.1–17) | 44 (37–53) | 3.0 | 2E-06 |
|  |  | *Cntn1* | 6.4 (4.9–8.5) | 14 (12–16) | 1.8 | 2E-02 |
|  |  | *Cxcr3* | 1.1 (0.8–1.6) | 5.5 (4.2–7.1) | 4.4 | 7E-04 |
|  |  | *Dbp* | 14 (8.4–23) | 81 (38–173) | 6.4 | 2E-03 |
|  |  | *Gzmk* | 1.2 (0.5–2.6) | 12 (8.2–17) | 4.8 | 4E-03 |
|  |  | *Hlf* | 4.1 (3.1–5.2) | 12 (8.5–18) | 2.9 | 1E-02 |
|  |  | *Lck* | 12 (10–15) | 26 (22–31) | 1.9 | 2E-03 |
|  |  | *Nr1d1* | 53 (37–77) | 141 (114–176) | 2.2 | 1E-02 |
|  |  | *Nr1d2* | 17 (13–22) | 35 (30–42) | 1.9 | 1E-02 |
|  |  | *Per3* | 2.4 (1.6–3.5) | 7.8 (5.0–12) | 3.0 | 3E-02 |
|  |  | *Plekha6* | 6.7 (4.7–9.5) | 15 (13–19) | 2.0 | 1E-02 |
|  |  | *Scn7a* | 16 (12–21) | 41 (323–51) | 2.2 | 4E-04 |
|  |  | *Scube1* | 5.1 (3.5–7.4) | 13 (11–15) | 2.0 | 6E-03 |
|  |  | *Septin1* | 15 (13–17) | 29 (24–35) | 1.8 | 6E-03 |
|  |  | *Sh2d2a* | 1.9 (1.3–2.6) | 6.2 (4.7–8.2) | 3.2 | 6E-03 |
|  |  | *Sox7* | 11 (7.3–16) | 24 (21–27) | 1.8 | 8E-03 |
|  |  | *Spns2* | 8.9 (6.8–12) | 19 (17–22) | 1.9 | 2E-03 |
|  |  | *Stac2* | 4.2 (2.5–7.0) | 14 (9.4–21) | 2.8 | 1E-02 |
|  |  | TCRA | 20 (17–24) | 48 (39–58) | 2.2 | 2E-05 |
|  |  | TCRB | 25 (21–31) | 54 (45–65) | 1.9 | 6E-03 |

\*Abbreviations: 95CI, 95% confidence interval; FC, fold-change group 2 vs group 1; FDR, false discovery rate; IGHV, immunoglobulin heavy chain variable region; TCR, T-cell receptor type A or B

Notable among up-regulated DEGs at day 3 were several interferon-stimulated genes (ISG), the protein products of which have virus restriction activity: *Apobec1, Ifi27l2a, Ifit1, Ifitm3, Isg15, Mx2, Rsad2*, and *Rtp4* (Figure 4A and Table 2 and Table S1). Other ISGs among the up-regulated DEGs were those for 2’-4’ oligoadenylate synthetase viral sensors *Oas1, Oas2, Oas3, Oasl1*, and *Oasl2*, and the interferon regulatory factors *Gbp4, Irf7*, and *Irf9*. The prominence on day 3 of virus-sensing, interferon, and anti-virus responses in the lung was also evident in the GO term analysis (Figure 4C). We also draw attention to two other DEGs at day 3: *Ido1*, a kynurenine pathway enzyme, which was reported as associated with anti-inflammatory responses in some patients with SARS-CoV-2 infection (42), and *Reg3g*, which encodes a C-type lectin with antibacterial activity.

In the lungs collected on day 6 there was still evident by DEG and GO term analyses a continuing response to virus (Figure 4B and 4D; Table S1). But also underway by that time was a cell-mediated response, including natural killer (NK) cell activity. Day 6 DEG products with these activities or associations included the cytotoxic T-cell marker *Cd8*, the cytotoxic effector *Prf1*, the granzymes *Gzma, Gzmb*, and *Gzmk*, the chemokine receptor *Cxcr3*, and the NK cell markers *Cd52, Klrk1*, and *Ncr1*. Similarly to what we had noted in *P. leucopus* after exposure to high dose lipopolysaccharide or a TLR2 agonist (39, 40), inflammation-associated genes that were distinguished here for their minimal to absent transcriptional response to the virus infection in the lungs were *Ifng*, encoding interferon-gamma, and *Nos2*, encoding nitric oxide synthase 2 (Table S1).

We followed up these observations from the genome-wide RNAseq of protein coding sequences with targeted analysis with the focus on two of the initial findings and of the differences between day 3 and day 6 among the virus-infected animals (Table S3). The first was of selected PRR and ISG genes (Figure 5). For both the PRR genes (*Cgas, Dhx58, Ifih1*), and Rigi and the ISG genes (*Apobec1, Gbp4, Isg15*, and *Mx2*) transcription was elevated over those of control animals at day 3 and had returned to close to control values by day 6. For all 8 genes, the two animals that had the highest sickness scores were not distinguishable from the other animals in their responses on day 3. These dynamics for the PRR and ISG genes corresponded to the dynamics for virus burdens in the lung for the infected animals (Figure 2).

**Figure 5.**
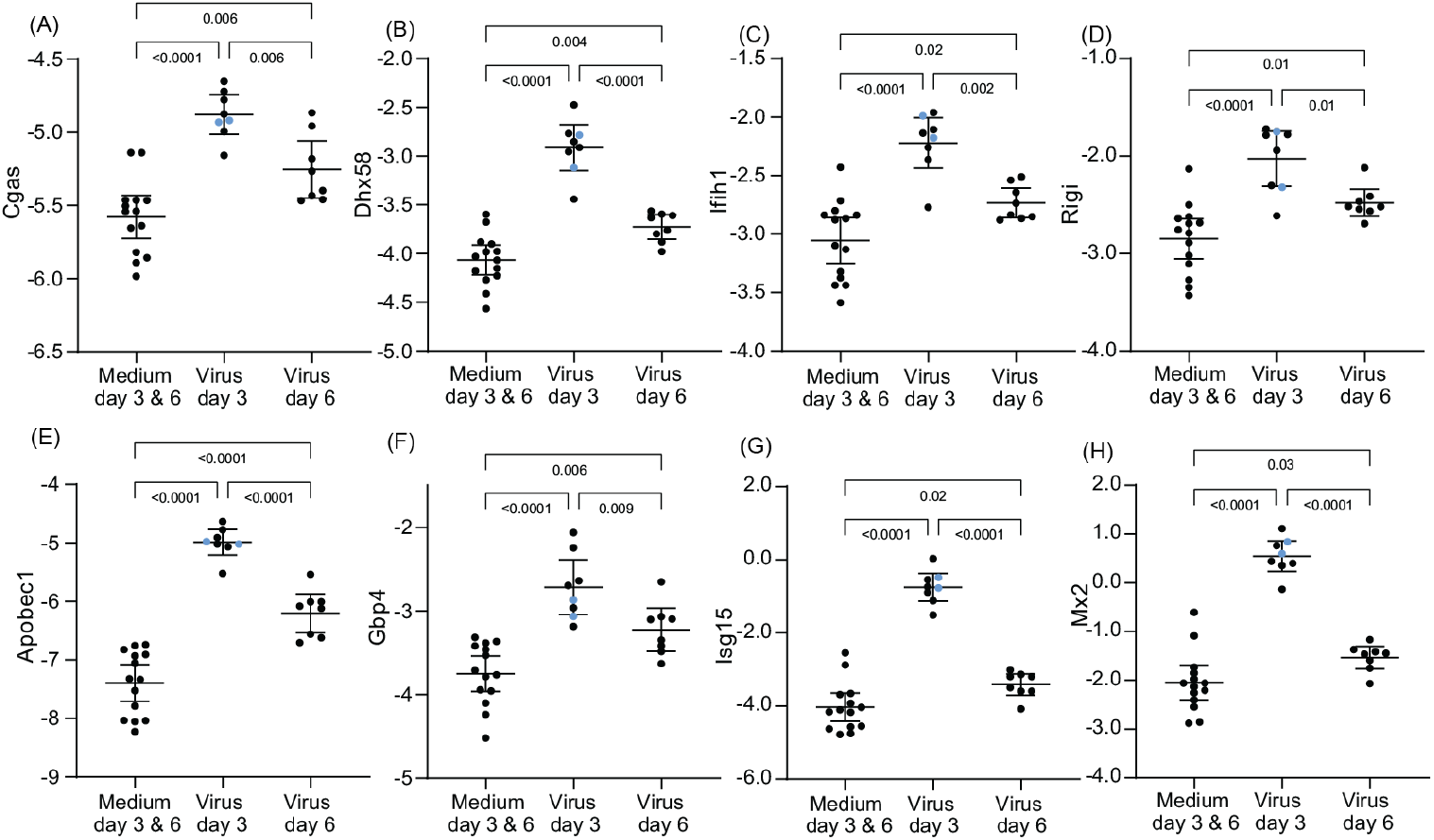
Differentially-expressed genes in lung tissue for select pattern recognition receptors (PRR) and interferon stimulated genes (ISG) of *P. leucopus* infected with SARS-CoV-2 virus or medium alone on days 3 or 6. Panels A-H are box-whisker plots with means and 95% confidence intervals of RNA-seq data as natural logarithms of target gene reads normalized for Gapdh transcription in the same sample (Table S3). The PRR genes of panels A-D are *Cgas, Dhx58, Ifih1*, and *Rigi*, and the ISG genes of panels E-H are *Apobec1, Gbp4, Isg15*, and *Mx2*. Each point represents an individual animal with two animals exhibiting severe disease with clinical scores of 3 (Table 1) indicated by blue dots. The displayed *p* values are from ANOVA.

In contrast to *Irf7*, which increased in transcription in infected animals on day 3 (Figure 4), *Irf3* was lower in transcription than control group values on that day before returning to a comparatively high baseline level (Figure S2). The two sickest animals were in the lower half of the distribution for *Irf3* on day 3 but in the higher half for *Irf7* on that day. A reciprocal relationship between these two transcription proteins has been characterized as a handoff from the constitutively-expressed *Irf3* to *Irf7* in a type 1 interferon pathway during viral infection (43). In this characteristic, *P. leucopus* was responding similarly to mice and humans, namely, *Irf3* activation driving *Irf7* and subsequently *Isg15* transcription (44).

Different transcription dynamics were observed among genes associated with cytotoxic T-cells and NK cells (Figure 6 and Table S3). Transcriptions were in general higher on day 6 than day 3, which in turn were higher than for control animals. This set included genes for immune cell markers and receptors, namely *Cd8, Cd52, Cxcr3, Ncr1* (natural cytotoxicity triggering receptor 1), *Tigit* (T cell immunoreceptor with Ig and ITIM domains), and *Tlrk1* (NKG2D), as well as cytotoxicity effectors *Gzmk* (granzyme K) and *Prf1* (perforin-1). For some of these genes, namely *Cd8, Cd52, Klrk1*, and *Prf1*, the two sickest animals had values in the lower halves of the distributions, suggesting an insufficiently developed cell-mediated response by day 3.

**Figure 6.**
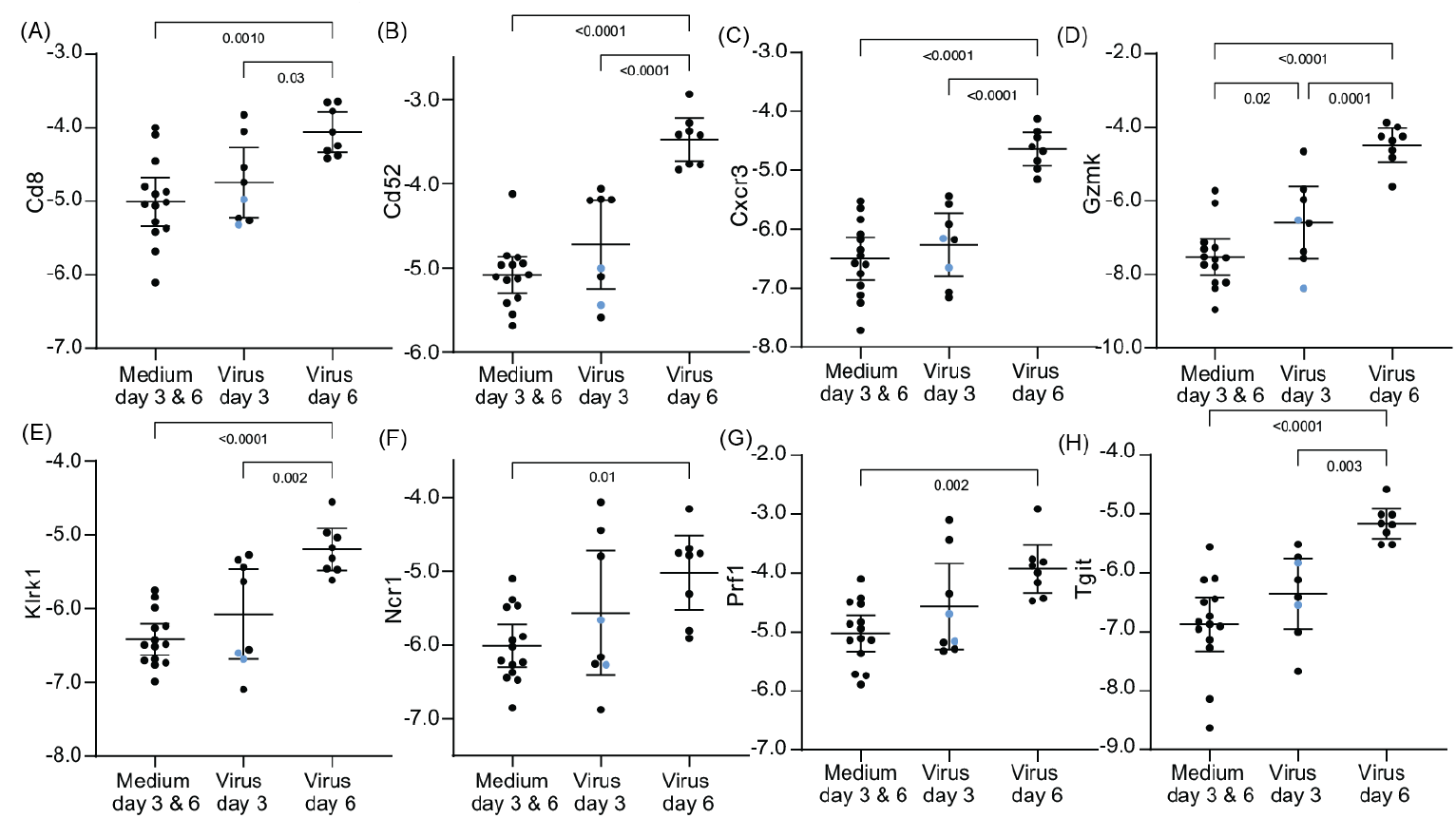
Differentially-expressed genes in lung tissue for select natural killer and cytotoxic T-cell markers of *P. leucopus* infected with SARS-CoV-2 virus or medium alone on days 3 or 6. Panels A-H are box-whisker plots with means and 95% confidence intervals of RNA-seq data as natural logarithms of target gene reads normalized for *Gapdh* transcription in the same sample (Tables S1 and Table 2). The genes are *Cd8, Cd52, Cxcr3, Gzmk, Klrk1, Ncr1, Prf1*, and *Tigit*. Each point represents an individual animal with two animals exhibiting severe disease with clinical scores of 3 (Table 1) indicated by blue dots. The displayed *p* values are from ANOVA.

### Comparison of *P. leucopus* to *M. musculus* in its response to the virus

For this study we did not carry-out a parallel experiment with *M. musculus* under the same conditions. So interpretations about the differences between species in their responses to SARS-CoV-2 infection are necessarily limited in scope. With this caveat in mind, we accessed a comparable RNA-seq data of lung from an earlier study by Winkler et al. of K18-hACE2 transgenic mice infected with the USA/WA-1 virus strain by the intranasal route at a comparable dose to that of the present study (45). In the mouse study samples were collected at day 2, 4, and 7, and uninfected mice served as controls. For the present study we combined the data for day 2 and day 4 time points and equated these with our day 3 time point and named both sets as “early”. Day 6 data of the present study and day 7 data from the mouse study were designated “late” sets for this analysis. We used the PE150 reads and metadata of that study that were publicly available The reference set for *M. musculus* was 22,762 non-redundant CDS sequence previously described (41). For the inter-species comparison of specific genes normalization of transcription was to the *Gapdh* gene of each species.

Figure 7A shows representative differences between the two species in log_2_ fold-changes to respective control animals under these particular circumstances. At the early time points in the infections there was higher transcription of ISGs, such as *Gbp4, Irf7, Isg15* and *Mx2*, in both species. At this stage three distinguishing genes were *Tnf* for tumor necrosis factor alpha, which was increased in transcription compared to controls in mice but not in the deermice, and *Apobec1* and *Slpi* (secretory leukocyte peptidase inhibitor), which were substantially increased in transcription in *P. leucopus*. Differences between species were more marked at the late time points. By that time the ISGs had decreased in transcription in *P. leucopus*, but these remained elevated, as did *Tnf*, in *M. musculus*. Two other stand-out differences at the late time points were elevated transcription of genes for interferon-gamma (*Ifng*) and nitric oxide synthase 2 (*Nos2*) in the *M. musculus* lungs but not in the lungs of infected deermice.

**Figure 7.**
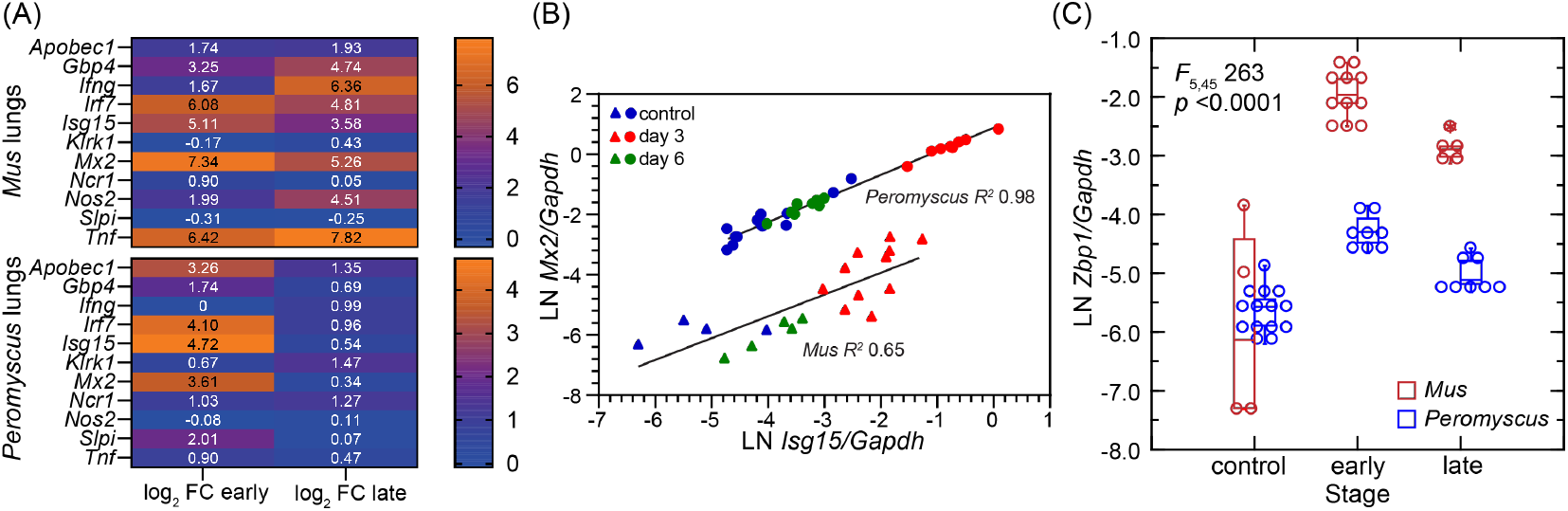
Cross-species comparison of transcriptional responses in lungs of *P. leucopus* and *M. musculus* species to SARS-CoV-2 infection. The data for *P. leucopus* are from Table S1. The data for the analysis for *M. musculus* are from our analysis (Table S4) of primary RNA-seq data from the study of Winkler et al. of K18-hACE2 mice (45). Combined data are at https://datadryad.org/dataset/doi:10.5061/dryad.cc2fqz6mm. The infections in the two species were denoted as “early” for day 3 for *P. leucopus* and combined day 2 and 4 for *M. musculus* and “late” for day 6 for *P. leucopus* and day 7 for *M. musculus*. <u>Panel A</u> shows heat maps for each species for genes *Apobec1, Gbp4, Ifng, Isg15, Klrk1, Mx2, Ncr1, Nos2, Slpi*, and *Tnf* with log_2_ values of fold-change (FC) differences between infected animals and uninfected controls in same periods. <u>Panel B</u> shows scatter plots with separate linear regressions and coefficients of determination (*R*^*2*^) for *Gapdh*-normalized *Mx2* transcription (*y*-axis) on *Isg15* transcription (*x*-axis). Circles represent values from *P. leucopus* and triangles from K18-hACE2 mice. Colors denote experimental groups: red for early infection, green for late infection, and blue for controls. <u>Panel C</u> has box-whisker plots with medians and 25th and 75th quartiles of *Gapdh*-normalized transcription of *Zbp1* for *P. leucopus* or *M. musculus* at different stages of infection. General Linear Model values are for stage nested within species.

While the antiviral effectors *Isg15* and *Mx2* elevated in the lungs by the early time points in both species, transcription of both genes was at a higher constitutive level relative to *Gapdh* transcription at baseline in *P. leucopus* (Figure 7B). In this characteristic of constitutive expression of certain ISGs, *P. leucopus* resembles some species of bats that serve as disease agent reservoirs (46). Another difference between species in this analysis was greater magnitude increase in infected mice in the transcription of *Zbp1*, the gene for the Z-nucleic acid sensing PRR Z-binding Protein 1 (Figure 7C).

### RNA-seq of the brain

The brain tissue samples from the controls and infected animals comprised the olfactory bulb as well as the cerebrum and cerebellum. For this tissue PE150 reads were reverse-transcribed from total RNA that had been first depleted of ribosomal RNA and then aligned with the full set of *P. leucopus* CDS (Table S5). This allowed quantitative estimation of viral RNA copies as well as mRNA and long non-coding RNAs in this tissue (Table 1). There was a high correlation between viral RNA copy numbers as measured by matching reads in bulk RNA-seq and as measured by RT-qPCR (Figure 8A). As with the RT-qPCR measures of viral RNA in the brain (Table 1), for the 8 virus-infected animals at day 3 or day 6 in each experiment, there was overall ∼100-fold more viral RNA reads (log_10_ mean [95% CI]) in brains of experiment 2 animals (3.5 [2.9-4.0]) than in experiment 1 (1.4 [0.9-2.0]) (*p* = 0.0002).

**Figure 8.**
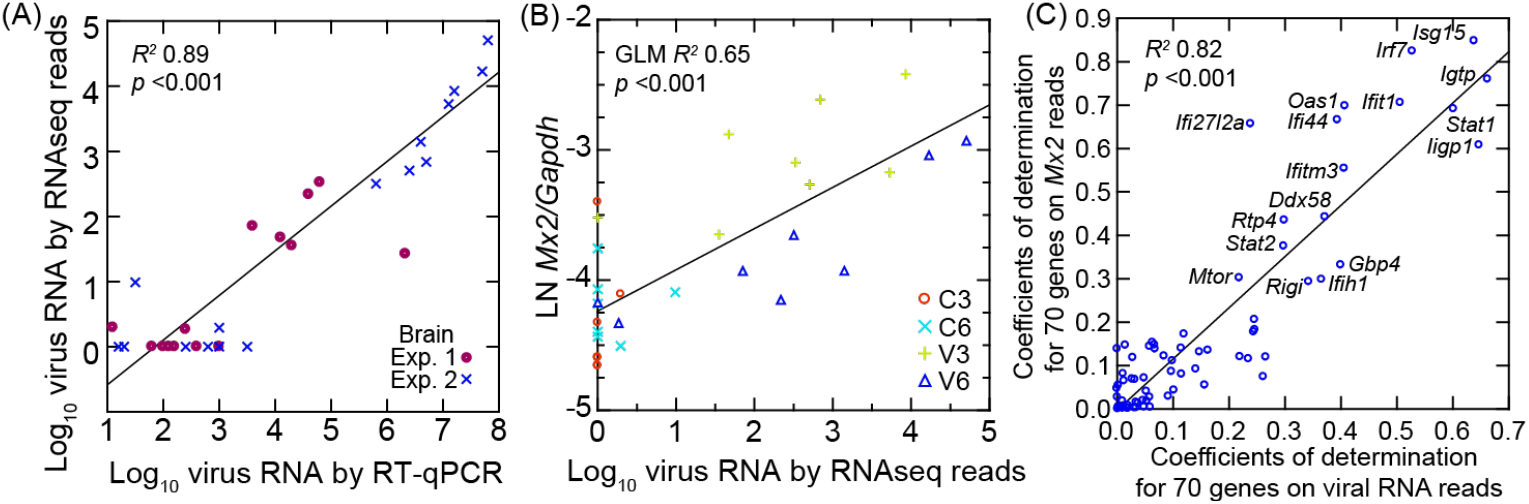
Scatter plots with regressions of analyses of SARS-CoV-2 virus in the brain of *P. leucopus* and selected genes for the host response. In all analyses cDNA libraries for genome wide and target RNAseq were from total RNA depleted of ribosomal RNA. In each panel the coefficients of determination (*R*^*2*^) for the regressions are shown. <u>Panel A</u> is linear regression of log-transformed viral RNA copies by RNAseq reads on viral RNA copies by specific RT-qPCR. The data points are distinguished by whether Experiment (Exp.) 1 or 2. The data for the graph is given in Table 1. <u>Panel B</u> is a General Linear Model (GLM) regression of natural logarithm (LN) of *Mx2* transcripts normalized for *Gapdh* on log-transformed viral RNA reads in the same RNAseq output. The second independent variable for the GLM was day of infection. <u>Panel C</u> is linear regression of coefficients of determination for each of 70 targeted genes with *Mx2* transcription on coefficients of determination for each of 70 targeted genes with viral RNA reads. The identities of genes with *R*^*2*^ of >0.3 with each of the two independent variables (with the exception of *Ifi27l2a*) are shown. The data for graphs are from Table S6.

By PE150 reads normalized for *Gapdh* transcription in the same samples, we also observed close associations between viral RNA levels and *Isg15*, with its transcription level at a similar viral load higher on day 3 than on day 6 (Figure 8B). From this observation we carried out targeted RNA-seq of 70 selected genes (Figure 8C and Table S6). Using these data we determined the coefficients of determination (*R*^*2*^) for each with either viral RNA reads for the sample or the normalized reads for the *Mx2* ISG for the sample. Plotting one of those series against the other revealed genes that very highly correlated with both viral RNA and *Mx2* transcription. In addition to some of the ISGs identified in the study of the lungs (Table 2), such as *Ifi27l2a, Irf7, Isg15, Rsad2, Rtp4*, and *Usp18*, we also observed elevated transcription in the virus-infected brain on day 3 of ISGs *Ifi35, Ifitm3*, and *Irgm2* (Table 3).

**Table 3.**
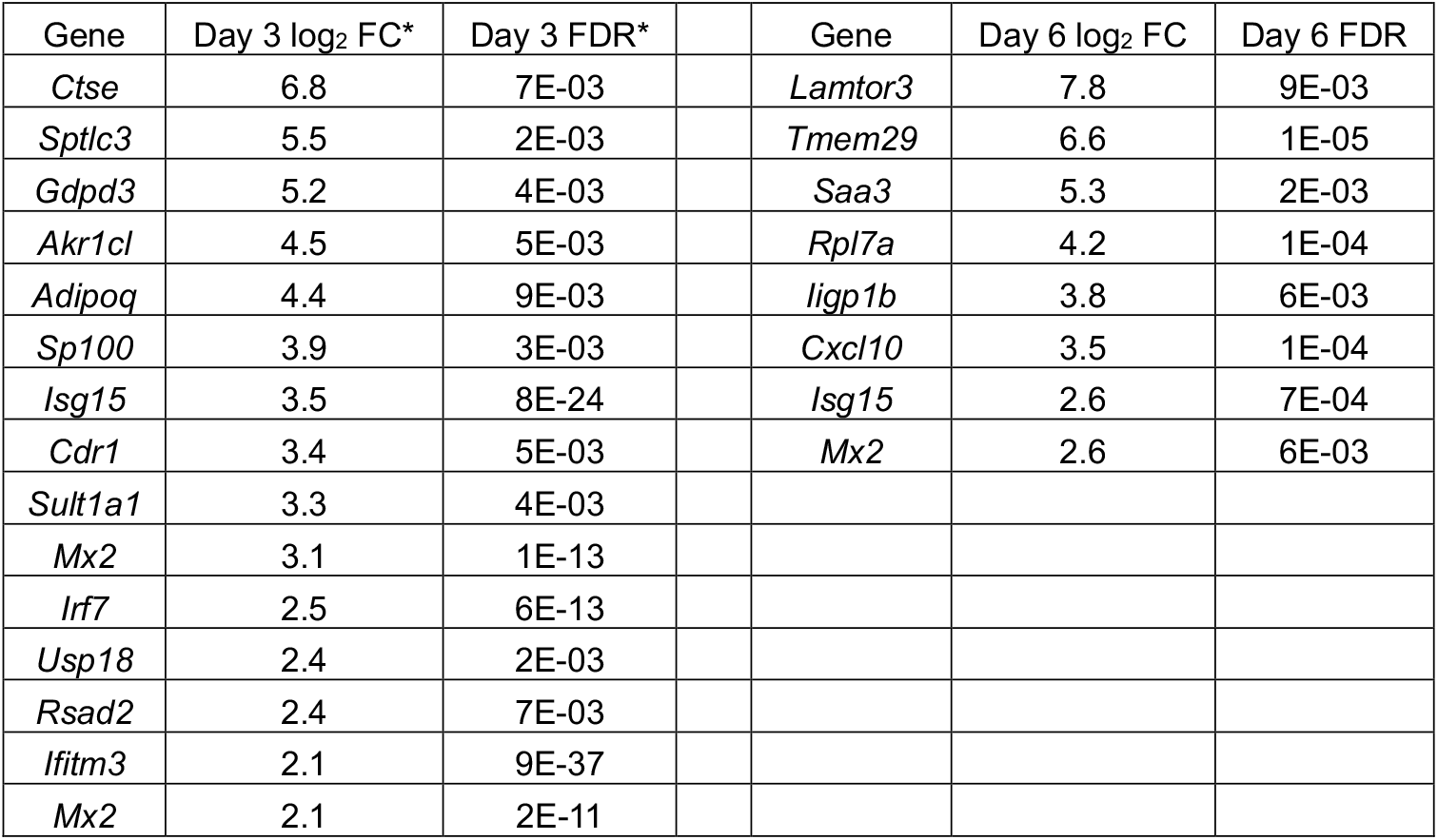
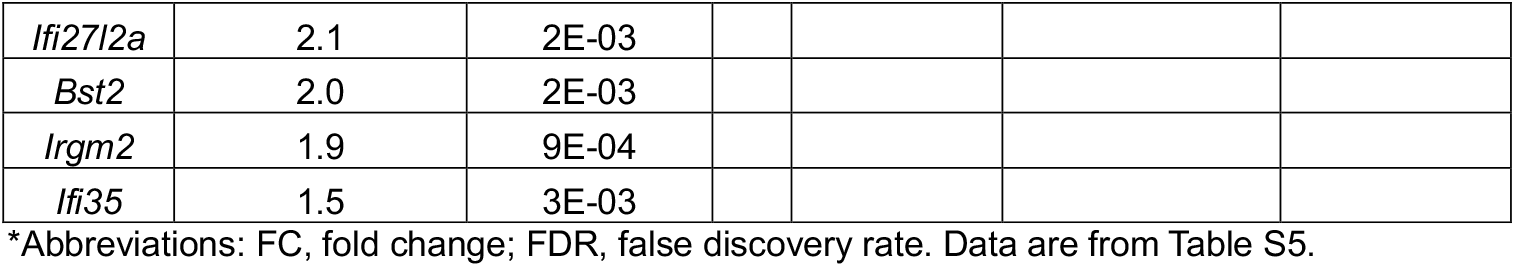
Up-regulated differentially-expressed genes in the brain on day 3 or day 6 of infection.

| Gene | Day 3 log<sub>2</sub> FC* | Day 3 FDR* | Gene | Day 6 log<sub>2</sub> FC | Day 6 FDR |
| --- | --- | --- | --- | --- | --- |
| *Ctse* | 6.8 | 7E-03 | *Lamtor3* | 7.8 | 9E-03 |
| *Sptlc3* | 5.5 | 2E-03 | *Tmem29* | 6.6 | 1E-05 |
| *Gdpd3* | 5.2 | 4E-03 | *Saa3* | 5.3 | 2E-03 |
| *Akr1cl* | 4.5 | 5E-03 | *Rpl7a* | 4.2 | 1E-04 |
| *Adipoq* | 4.4 | 9E-03 | *ligp1b* | 3.8 | 6E-03 |
| *Sp100* | 3.9 | 3E-03 | *Cxcl10* | 3.5 | 1E-04 |
| *Isg15* | 3.5 | 8E-24 | *Isg15* | 2.6 | 7E-04 |
| *Cdr1* | 3.4 | 5E-03 | *Mx2* | 2.6 | 6E-03 |
| *Sult1a1* | 3.3 | 4E-03 |  |  |  |
| *Mx2* | 3.1 | 1E-13 |  |  |  |
| *Irf7* | 2.5 | 6E-13 |  |  |  |
| *Usp18* | 2.4 | 2E-03 |  |  |  |
| *Rsad2* | 2.4 | 7E-03 |  |  |  |
| *Ifitm3* | 2.1 | 9E-37 |  |  |  |
| *Mx2* | 2.1 | 2E-11 |  |  |  |

|  |  |  |
| --- | --- | --- |
| *Ifi2712a* | 2.1 | 2E-03 |
| *Bst2* | 2.0 | 2E-03 |
| *Irgm2* | 1.9 | 9E-04 |
| *Ifi35* | 1.5 | 3E-03 |

\*Abbreviations: FC, fold change; FDR, false discovery rate. Data are from Table S5.

The gene for the chemoattractant CXCL10, a chemokine also known as interferon-gamma inducible protein 10, was notable for its higher level of transcription in the brain of infected animals on day 6 than on day 3, a result confirmed by RT-qPCR (Figure 9, Table 3 and Table S6). This was in contrast to the lungs where *Cxcl10* transcription was marginally higher on day 3 than on day 6 (Table S1). *Isg15* and *Mx2* were two ISGgenes that remained elevated in transcription in the brain on day 6 (Table 3).

**Figure 9.**
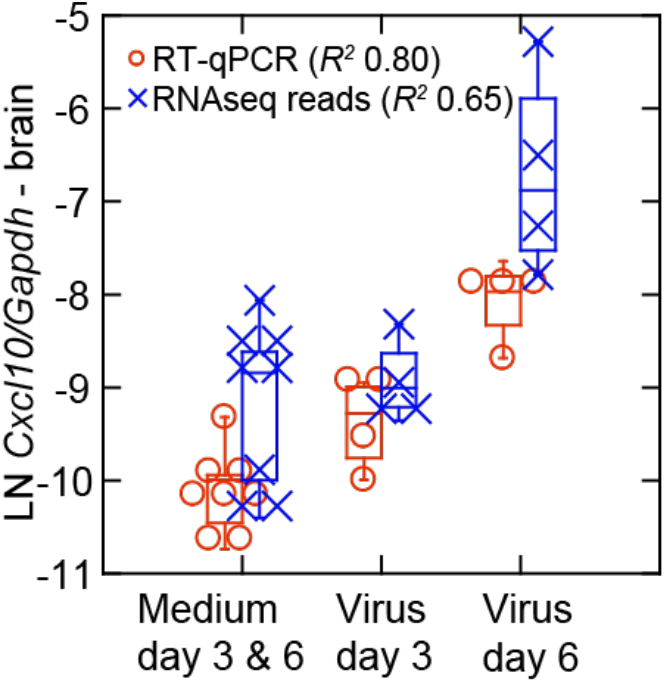
Late transcriptional response for the chemokine Cxcl10 in the brain of *P. leucopus* infected with SARS-CoV-2 virus for experiment 2. The box-whisker plot showing the median and 25th and 75th quartiles is of transcriptions of *Cxcl10* normalized for *Gapdh* in brain tissue extracts from controls (medium alone) or virus-infected animals on either day 3 or day 6 of infection. Transcription was assessed by targeted RNA-seq and by RT-qPCR of the same RNA extract for Cxcl10 and Gapdh transcripts (Table S5). Coefficients of determination (*R*^*2*^) were determined for each analysis.

## Discussion

Using another species as an example, we confirmed that deermice of the genus *Peromyscus* can be infected by the respiratory route with the SARS-CoV-2 coronavirus and, further, that this is followed by dissemination to the brain and gastrointestinal tract. The susceptibility finding was not unexpected, given the reports of successful infections among *P. maniculatus* sub-species and *P. californicus* (31-33). The strain of virus and the approximate inoculum were the same as studies of Griffin et al. and Lewis et al. (31, 33). Besides providing a fuller characterization and development of the *Peromyscus* model, the study assessed the extent to which *P. leucopus* could both keep the virus in check and forstall pathology from the host response. We knew of the white-footed deermouse’s capacities on both those fronts for the several pathogens, such as *B. burgdorferi*, for which it serves as a competent reservoir. But what about a potentially lethal virus that from the serological evidence it had not encountered before (34, 35)?

Each of the 16 animals that were inoculated with virus became infected by the criteria of an antibody response to the S antigen by day 6 and the presence of virus RNA in lung and/or brain by RT-qPCR (Table 1). These were at levels that could not be attributed to the inoculum alone without replication of the virus. In representative cases virus was recovered from lung or directly detected by hybridization. Even when detectable viral RNA was present in the lung and brain, 14 of 16 infected animals showed either mild illness or no overt signs of sickness at either day 3 or day 6, the termination points in both experiments. The two animals that were more demonstrably sick were male and between 919 and 932 days of age, a finding that calls to mind the increased risk of more severe disease observed in human studies for older males infected with SARS-CoV-2 virus (47). On the other hand, there were two other males of similar age who manifested only mild illness in experiment 2.

While the same strain of virus at the same titer were the inocula in each experiment, the greater degree of sickness recorded in experiment 2, along with generally higher viral RNA copies in the lung and brain in the second experiment, suggest differences in either infectiousness or virulence between the two virus batches, which came from different sources. This is an acknowledged limitation of the study. But accepting this constraint, one can interpret the differences between inocula between the two experiments as different dosages of the virus. From that perspective, the animals receiving the preparation representing a higher effective dose of infectious virions manifested more sickness in general and higher titers of virus in the brain (Table 1). On the other hand, as these were outbred animals sampled from a heterogeneous stock, the more severe disease in two animals might also be attributable to genetic variation between individual animals in responses to infection, as occurs in humans (48).

There were pathological changes in the lungs of infected animals that exceeded in degree the mild inflammation noted in controls that inhaled medium without virus. The findings in the lungs of infected *P. leucopus* were similar to what was observed for two subspecies of *P. maniculatus* (31, 32). In the study of Lewis et al. of two other subspecies of *P. maniculatus*, as well as *P. polionotus* and *P. californicus*, among all the animals under study only a minority of the *P. californicus* animals in the study became severely ill, similar to what we observed here for *P. leucopus* (33). In the case of *P. californicus* more severe disease may have been associated with the hepatic steatosis that can occur in this species (49). Liver disease has been another risk factor for a higher case fatality among humans with SARS-CoV-2 infection (50). By gross examination of the organs there was no evidence of liver abnormalities in the two sickest *P. leucopus* at termination, but we cannot exclude liver dysfunction as an alternative explanation.

Across both experiments in the lungs of infected animals there was a biphasic immune response by genome-wide and targeted RNA-seq that has been noted in experimental infections of other mammals with SARS-CoV-2 (51, 52). In general, on day 3 the response was dominated by apparent up-regulation of genes for cytoplasmic PRRs and ISGs of the sort that would be expected in the early stage of viral infection (Figures 4 and 5). Besides such ISGs as *Isg15* and *Mx2*, we noted the marked elevation of transcription of *Apobec1*. The APOBEC family of cytidine deaminase enzymes are antiviral effectors that restrict replication through hypermutation and degradation of viral genomes (53, 54). Among the day 3 DEGs, another ISG of less reknown was *Ifi27l2a* (interferon, alpha-inducible protein 27 like 2A), also known as *Isg12* and a member of the IFI6/IFI27 protein family. The human homolog IFI27 was reported to inhibit activation of the PRR MDA5 (Ifih1), thereby countering that particular pathway in response to SARS-CoV-2 (55). Elevated transcription on day 3 of *Isg15* in the lungs, along with *Oas2* and *Cxcl10*, was also noted by Griffin et al. (31).

By day 6 most of these DEGs were reduced in transcription if not back to level of controls. Instead, we observed by that time evidence of the cell-mediated immune response with prominence for cytotoxic T cell and NK activity, as evident in the heightening transcription over days 3 and 6 of the cytotoxic effectors granzyme K and perforin-1 (Figures 4 and 6). While a mild late increase in markers and effectors associated with cytotoxic T cells and NK cells was also observed in the secondary analysis of *M. musculus* infected with the virus, among the infected mice there was not a substantial decline by that later time point in the transcription of the ISG genes *Apobec1, Gbp4, Isg15*, and *Mx2*, or that of the *Zbp1* gene for the Z-nucleic acid sensor (56) (Figure 7). The SARS-CoV-2 virus, among some other viruses, activates an inflammatory cell death pathway through ZBP1 sensing (57). A higher initial expression of *Abobec1* in *P. leucopus* may have had a restraining effect on the ZBP1-mediated inflammation (58).

Three genes for pro-inflammatory proteins that had risen in expression in the *M. musculus* by the second time point were *Ifng, Nos2*, and *Tnf*. The two experimental models are not fully commensurate because of the more severe disease in the transgenic *M. musculus*, but the findings of comparatively low expression of *Ifng* and *Nos2* in deermice are consistent with what we observed for *P. leucopus* and *M. musculus* in the LPS models of sepsis (39, 40). Griffin et al. also reported that transcription of *Ifng* at low levels was noted only in a subset of infected *P. maniculatus* (31).

Another similarity between the interspecies comparison for this study and our previous studies of the blood of animals treated with LPS (39, 40), as well as the study of primary dermal fibroblasts treated with the TLR2-agonist lipopeptide Pam3CSK (41), was the higher baseline transcription on a normalized basis of the ISGs *Isg15* and *Mx2* in deermice or their cells than mice samples under the same conditions. While the full implications of these host distinctions await results of future investigations, a plausible inference at this point is that higher constitutive expression of antiviral effectors like *Isg15* and *Mx2* provide a leg-up for the animals at the time of exposure to a new virus.

Interpretations of the results for the brain are restricted in scope for the following reasons: (a) There apparently was less virus in the brain in experiment 1 than in experiment 2, as we discuss above. (b) The brain tissue included the olfactory bulb as well as the cerebrum, brain stem, and cerebellum. While the olfactory bulb is part of the brain, the route for its infection can include transmission through the cribiform plate of the skull from the nasopharynx as well as from dissemination through the circulation (59). Accepting these limitations, inferences of informative value are possible if one uses viral RNA copies in the brain as the independent variable (Figure 8). There was no evident pathology by microscopy, but the RNA-seq results were consistent with a response to viral infection on both day 3 and day 6 (Figure S3). The finding is consistent with a low level of viral presence in the brain and a host response detectable at the transcriptional level without accompanying overt histopathological changes. This was evident from the elevations ISGs, including *Ifi27l2a*, transcription of which is associated with responses in the central nervous system (60). Another notable finding in the brain samples was the transcription of the chemokine *Cxcl10*, which was increasing through day 3 and day 6 (Figure 9). While CXCL10 has been associated with pro-inflammatory responses, a more definitive test of its role using a *Cxcl10* knock-out mouse and a mouse-adapted strain of the SARS-CoV-2 virus indicated that CXCL10 provides protection against mortality from this virus (61).

After review of the findings, can one conclude that *P. leucopus* “tolerated” infection with the SARS-CoV-2 virus, a pathogen, with which this species, by the evidence to date, was unlikely to have had prior contact? In the case of the 14 of 16 animals which manifested only mild or no sickness behavior or distress while experiencing extensive pathological changes in the lung, the answer is yes. By day 6, the animals still in the experiment were in recovery. Another indication of infection tolerance and moderation of inflammation in *P. leucopus* were undetectable to very low transcription of genes for interferon-gamma and nitric oxide synthase 2 in lungs, which contrasted with what was observed in *M. musculus* (Figure 7). The findings indicate that the trait of infection tolerance accommodates to pathogens that have not been encountered before, not only within a sample of the existing population but likely among their ancestors as well.

## Methods

### Animals

The *P. leucopus* was the LL stock of the Peromyscus Genetic Stock Center of the University of South Carolina (5). Animals were maintained on a 12 hr light/dark cycle in temperature- and humidity-controlled rooms, with ad libitum access to water and 8604 Teklad Rodent Diet of Harlan Laboratories (USA). Animal procedures were conducted under protocols AUP-18-020 and AUP-21-007 approved by the Institutional Animal Care and Use Committee of the University of California, Irvine and performed in accordance with Guide for the Care and Use of Laboratory Animals: Eighth Edition (National Academies Press).

### Virus

The SARS-CoV-2 virus strain was 2019-nCoV/USA-WA1 from BEI Resources (USA) catalog NR-52281 for experiment 1 and Microbiologics (USA) catalog G2027B for experiment 2. Virus was propagated in Vero E6 cells from ATCC (USA) catalog CRL-1586 in Dulbecco’s Modified Eagle Medium (DMEM) from Sigma-Aldrich (USA) supplemented with 25 mM glucose, 1% HEPES, and 2% fetal bovine serum (FBS) at 37°C with 5% CO_2_. Virus titers as plaque forming units (pfu) were determined by focus forming assay (see below) in Vero WHO cells from ATCC catalog CCL-81, as described by Case et al. (62) for experiment 1 or in Vero E6 cells, as described by Prakash et al. (63) for experiment 2.

Focus forming assay for virus titer. Lung tissues in phosphate-buffered saline were homogenized in a Closed Tissue Grinder System from Fisherbrand (USA). Homogenates were centrifuged at 1000 x g for 15 s to pellet debris. The supernatants were stored at -80°C. For the assays performed in duplicate an aliquot of supernatant was thawed and then applied to Vero E6 cells in a 96-well plate at a density of 5 × 10^4^ cells/well in DMEM with 10% FBS and then overlaid with 1% (w/v) methylcellulose (Sigma-Aldrich). The cultures were incubated at 37°C for 1 h with gentle rocking of the plate every 15 min and then an equal volume of 2% methylcellulose was added to each well. The plates were incubated at 37°C and 5% CO2 for 24 h and then fixed with 10% neutral buffered formalin. Infected foci were detected and quantitated by incubation at 4°C first with rabbit anti-SARS-CoV-2 nucleocapsid antiserum from Novus Biologicals (USA) catalog NB100-56576) and then, after washing, with horseradish peroxidase (HRP)-conjugated anti-rabbit IgG secondary antibody of BioLegend (USA). Signal was developed with True Blue HRP substrate (Sigma-Aldrich) and measured on an iSpot ELISpot instrument of Autoimmun Diagnostika (Germany).

### Experimental infections

Animals were transferred to an Animal Biosafety Level 3 (ABSL-3) facility at University of California Irvine at least 24 h prior to inoculation. Animals in the ABSL-3 were housed individually in IsoCage system cages of Techniplast USA (USA) under negative pressure. Control animals were housed under the same conditions in an ABSL-2 facility. The final titers of the virus stock were ∼6 × 10^6^ pfu/ml for each experiment. Equal numbers of females and males were used for each group and condition. For experiments 1 and 2 on day 0 animals were weighed and were infected intranasally. An animal was lightly anesthetized with isoflurane, held upright, and a 20 µl volume of medium alone (controls) or with an estimated 2 × 10^4^ virus particles was applied dropwise to the nares. Animals were held in position until they inhaled the deposited fluid as they wakened. Animals were returned to their cages and then monitored daily in the ABSL-3 facility for distress and sickness behavior. Scoring for this was 0, 1, 2, or 3 according to the following criteria: 0 for no visible evidence of distress or sickness behavior throughout the experiment, 1 for ruffled fur (piloerection) but active and alert, 2 for ruffled fur and mild-moderate lethargy but rousable, and 3 for ruffled fur, marked lethargy, tachypnea, and reduced mobility when handled. Control and virus-inoculated animals were euthanized on either day 3 or day 6 by exposure to high carbon dioxide partial pressure. After cessation of breathing, the chest cavity was opened; blood was collected by cardiac puncture and transferred to lithium heparin tubes from Becton Dickinson (USA). The anticoagulated blood was centrifuged at 7,000 x g for 3 min. The plasma was heat-inactivated at 56°C for 45 min and then frozen at -80°C until use. After lungs were excised, halves of right lungs were homogenized in TRI Reagent from Zymo Research (USA). Freshly-obtained right-side brain tissues were frozen in Invitrogen TRIzol reagent of Thermo Fisher Scientific (USA) and stored at -80°C. The remaining tissue of the right lung was snap-frozen on dry ice and stored at -80°C. Left lungs or left-side brain tissues were immediately fixed after dissection in 10% neutral-buffered formalin for 24 h at room temperature and then placed in 70% (v/v) ethanol.

### Histopathology

Fixed tissues were paraffin-embedded, sectioned, mounted on glass slides, and stained to hematoxylin and eosin. Slides were examined with a Leica DM1000 microscope equipped with a Leica ICC50 digital camera. At least 20 random high-power fields were evaluated for slide. Pathologic changes in lung tissues were characterized by type and severity using the criteria and scoring system of Matute-Bello et al. for experimental acute lung injury in animals (64). For each field, a score of 0, 1, or 2 was assigned based on the following parameters: (A) neutrophils in the alveolar space (0 =none, 1 = 1-5 cells, and 2 = >5 cells; (B) neutrophils in the interstitial space (0 = none, 1 = 1-5 cells, and 2 = >5 cells; (C) hyaline membranes (0 = none, 1 = 1 membrane, and 2 = >1 membrane); (D) proteinaceous fluid and cellular debris in alveoli (0 = none, 1 = present in one alveolus, and 2 = >1 alveolus); and (E) alveolar septal thickening (0 = septal thickness < 2x that of normal lung; 1 = 2x-4x thickening; and 2 = >4x thickening). The overall lung injury score for each specimen was calculated using the formula: [(20 x A)+ (14 x B) + (7 x C) + (7 x D) + (2 x E)]/(number of fields x 100).

### In situ hybridization

Slide-mounted sections of formalin-fixed, paraffin-embedded lung tissue from infected or uninfected *P. leucopus* were deparafinized, treated with hydrogen peroxide, incubated with protease, hybridized with probe, subjected to amplification, and then counterstained with hematoxylin according the manufacturer’s instructions for the RED RNAscopeTM 2.5 HD Detection Kit of Advanced Cell Diagnostics (USA). The RNAscope probes used were the following: V-nCoV2019-S (catalog 848561) for the SARS-CoV-2 virus (positions 21631-23303 of NC-045512), Hs-PPIB (catalog 313901; peptidylprolyl isomerase B gene) as positive control for lung tissue, and *Bacillus subtilis* DapB (catalog 310043; positions 414-862 of EF191515) as a negative control. As a positive control for the SARS-CoV-2 virus brain tissue of infected K18-hACE2 *Mus musculus* was used (65). Stained tissues were viewed at 400x magnification with an Olympus BX60 microscope equipped with a Nikon DS-Fi3 model digital camera.

### Recombinant SARS-CoV-2 protein

HEK293T cells from ATCC catalog CRL-3216 in 143 cm^2^ tissue culture dishes of Genesee Scientific (USA) with DMEM with 10% FBS and 20 mM glutamine. After 24 h the cultured cells were washed twice with Dulbecco’s Phosphate Buffered Saline (DPBS) and suspended in BalanCD HEK293 serum-free medium from Irvine Scientific (USA). Cells were transfected with pPP14 SARS-CoV-2 NFL 2P Foldon-His plasmid, which encodes His-tagged spike protein and was provided by Rogier Sanders (Weill Medical College of Cornell University) (29). Enrichment and purification of His-tagged protein in the culture supernatant was achieved by differential filtration and centrifugation and by affinity chromatography for the His tag. Protein concentration was determined at 280 nm on a NanoDrop spectrophotometer (Thermo Fisher Scientific). Purity of the recombinant protein was assessed as 95% by SDS-PAGE followed by staining with Simple Blue (Thermo Fisher Scientific). Recombinant nucleocapsid N protein was obtained from SinoBiological (USA) catalog 40588-v08b.

### Enzyme-linked immunosorbent assays (ELISA)

Flat bottoms of 96-well ELISA plates (Corning; 3690) were coated overnight at 4°C with either recombinant S protein (100 ng per well) or nucleocapsid (N) protein (50 ng per well) in DPBS. Wells were washed 3 times with DPBS containing 0.5% Tween-20 (wash buffer) and blocked for 1 h at 37°C with wash buffer with 5% non-fat dry milk (blocking buffer). Following 2 washes, plates were incubated for 1 h at 37°C with 3-fold serial dilutions of P. leucopus plasma in blocking buffer. Wells were washed 3 times with and then incubated for 1 h at 37°C with HRP-conjugated goat anti-*Peromyscus* IgG from SeraCare (USA) diluted 1:1000 in blocking buffer. Wells were washed 4 times, and the signal was developed after the final wash by addition of 50 µl of 1-Step TMB substrate (Thermo Fisher Scientific) to each well. Development was stopped by addition of 50 µl 1M H2SO4. The positive antibody controls were human monoclonal antibodies COVA1-18 for S protein and COVA103-C12 for N protein and provided by Marit van Gils (Amsterdam University, Netherlands) (29). Absorbance was measured at 450 nm on a BioTek ELx808 plate reader. The endpoint titer was defined as the highest plasma dilution yielding an absorbance value that exceeded the mean of 16 blank wells plus 3 standard deviations. Assays were performed in duplicate.

### Neutralization assay with pseudotyped virus

HEK293T cells were co-transfected in presence of polyethylenimine with HIV-1 NL4-3 Gag-iGFP ΔEnv plasmid (NIH AIDS Reagent Program, National Institute of Allergy and Infectious Diseases, USA) (66) and the SARS-CoV-2 spike-expressing plasmid pcDNA3.1 SARS-CoV-2 S, which was a gift of Thomas Gallagher of Loyola University (USA) (67). After 3 d culture supernatants were harvested, clarified by centrifugation, aliquoted, and stored at -80°C. For the neutralization assay, 96-well flat bottom tissue culture plates (Genessee Scientific) for incubation for 1 h at 37°C of 25 µl pseudotyped virion suspension and 25 µl of heat-inactivated *P. leucopus* plasma serially diluted 1:30 fold in DMEM/FBS starting at 1:100. Then 2.5 × 104 HEK293T cells stably expressing human ACE2 (BEI Resources) were added to each well 50 µl volumes, and the plates were incubated for 48 h at 37°C. Culture supernatants were removed, and cells were detached from each well by addition of 100 µl Accutase cell detachment solution from Innovative Cell Technologies (USA). The suspended cells were transferred in 100 µl volumes to round-bottom 96-well plates (Genesee Scientific), washed once with DPBS, and fixed with 10% formalin at 4°C for 30 min. Measurement of fluorescence of the Green Fluorescent Protein expressed from the plasmid in the fixed cells was done on a NovoCyte (USA) flow cytometer with excitation at 488 nm and emission measured at 519 nm. The amount of neutralization was estimated by comparing the median median fluorescence intensity of test wells to wells without antibody exposure. The 50% inhibitory dilution (ID50) titers in duplicated were calculated as the plasma dilution that reduced fluorescence by 50%. The positive and negative control antibodies for the assay were aforementioned monoclonal antibodies COVA1-18 and COVA103-C12, respectively.

### RNA extraction

Lung RNA was isolated from the homogenized tissue suspension using the Direct-zol RNA Miniprep Kit (Zymo Research), following the manufacturer’s protocol. RNA was extracted from brain tissue in TRIzol that had been stored at −80°C by first mechanically homogenizing in a TissueLyser of QIAGEN (USA) apparatus with 3-mm stainless steel beads. This was followed by chloroform phase separation, ethanol precipitation, DNase I treatment, and finally purification with the RNeasy Mini Kit (QIAGEN). RNA concentrations were determined using both a NanoDrop spectrophotometer and a Qubit fluorometer (Thermo Fisher Scientific). RNA quality was assessed using an Agilent (USA) 2100 Bioanalyzer with the Nano RNA chip. All RNA was stored in RNase-free distilled water at -80°C.

### RT-qPCR of virus RNA

For each 25 µl reaction volume, 500 ng of RNA was used as template with 0.1 µM each of SARS-CoV-2 nucleocapsid gene–specific primers (forward: 5′-GGGGAACTTCTCCTGCTAGAAT-3′; reverse: 5′-CAGACATTTTGCTCTCAAGCTG-3′) and the qPCRBIO SyGreen 1-step Go Hi-ROX kit of PCRBiosystems (UK). Amplification was performed using a Rotor-Gene 6000 real-time PCR system (QIAGEN) with the following cycling conditions: 50°C for 30 min, 95°C for 15 min, followed by 45 cycles of 94°C for 15 s and 60°C for 20 s. Quantitative virus copy numbers in the samples were estimated in triplicate using a standard curve generated from serial dilutions of virus RNA, measured RNA concentrations, and molecular weight of a virus genome.

### Sequencing

For lung samples production of cDNA libraries was with TruSeq Stranded mRNA kit of Illumina (USA) and for brain samples production of cDNA libraries with Illumina TruSeq Stranded Total RNA kit with ribosomal RNA depletion. After normalization and multiplexing, the libraries were sequenced on an Illumina NovaSeq 6000 instrument at the U.C. Irvine Genomics High-Throughput Facility. There were 100 cycles of paired-end chemistry reads (PE100) targeting 80 million reads per sample for lung, and a targeted 40 million PE150 reads for brain. The quality of sequencing reads was assessed using FastQC of Babraham Bioinformatics (USA). Adapter sequences, homopolymeric 5’ or 3’ ends, and low-quality reads (Phred score <15) were trimmed using Trimmomatic (68).

### Third party RNA sequence data for *Mus musculus*

Publicly-available fastq format files of sequencing reads of lung samples from a study of experimental infection in K18-hACE2 transgenic mouse model were accessed for comparisons of SARS-CoV-2 lung infections in *P. leucopus* with those reported for *M. musculus* (45). The BioProject was PRJNA64513, and the corresponding BioSamples were SAMN15493425-15493448. National Center for Biotechnology Information (NCBI) SRA files SRR12185412-12185431 comprised paired-end reads after 150 cycles Illumina reads from a mRNA library. The mean number (and range) of paired-end 150-bp reads for 20 samples were 7.9 (6.5-9.4) x 10^7^.

### RNA-seq analysis

Batched fastq files were subjected to RNA-seq analysis with CLC Genomics Workbench version 24 (QIAGEN). For the analysis of *P. leucopus* reads the source for the 22,654 protein coding sequences (CDS) of the genome of a *P. leucopus* LL stock female was GenBank accession number GCF_004664715.2 (36). Additional annotation, which was carried out manually using the default settings for the blastx algorithm (https://blast.ncbi.nlm.nih.gov) against non-redundant *M. musculus* protein sequences in GenBank, assigned gene names to ∼1000 sequences that had originally been assigned “LOC1146xxxxx” names by the NCBI vertebrate genome annotation pipeline. This was done iteratively as “LOC” sequences were identified as candidates for differentially expressed genes. The database with newly assigned gene names as well as original “LOC” designations and GenBank accession numbers for the protein products is available as “Table CDS_Pleuc_names_v7” at the Dryad Digital Repository (https://doi.org/10.5061.dryad.fqz612k44). For the *P. leucopus* data, the settings for length fraction, similarity fraction, and mismatch/insertion/deletion cost parameters were 90%, 0.4, and 3, respectively, for lung reads; for brain reads the length fraction was reduced to 0.35. The third-party *M. musculus* PE150 reads for lung were mapped against a reference set of 22,761 non-redundant protein-coding sequences derived from the strain C57BL/6 *M. musculus* reference genome (GCF_000001635.27_GRCm39) using the same parameter settings as for *P. leucopu*s lung reads.

### Genome-wide differential gene expression

Differential gene expression analysis was performed with *R* v. 4.4.3 with RStudio v. 2024.12.1 (69) as the front-end and the edgeR package v3.42.4 (70). Transcripts per million (TPM) from RNA-seq were used as input for normalization and dispersion estimation. edgeR was applied to TPM-normalized values rather than count data, which can affect dispersion and FDR estimates; counts-based models were further used to confirm effect sizes and the number of DEGs. Differential expression between experimental conditions was calculated using generalized linear models. Fold changes between two conditions were log_2_-transformed for graphics visualization. Transformed ratios were used for calculations of means and 95% confidence intervals (CI) and converted back as anti-logs for asymmetric 95% confidence intervals. The *p*-values were corrected for multiple testing using Benjamini-Hochberg method for false discovery rate (FDR) (71).

### Gene ontology term analysis

Differentially expressed genes identified by edgeR analysis and meeting the criteria of FDR <0.05 and log_2_ fold change >1.5 were selected for GO term enrichment analysis for both lung and brain datasets. Functional annotation and pathway enrichment were performed using Metascape (https://metascape.org) (72). Enrichment analysis was first conducted using the hypergeometric test. Similarity matrices were hierarchically clustered, and a 0.3 similarity threshold was used to define discrete functional clusters. Terms with the lowest *p*-value in each cluster were retained for representation in horizontal bar plots. GO terms refer to the Gene Ontology resource (http://geneontology.org), while additional annotations were derived from the KEGG Pathway (https://www.kegg.jp, denoted ‘mmu.’), WikiPathways (https://www.wikipathways.org, ‘WP…’), and Reactome (https://reactome.org, ‘R-MMU…’) databases.

### Targeted RNA-seq analysis

Targeted RNA-seq analyses of selected CDS from the genome-wide sets were performed using CLC Genomics Workbench. For the brain samples an additional reference was the complete sequence of the SARS-CoV-2 virus strain USA-WA1/2020 (GenBank accession MN985325). Paired-end reads for *P. leucopus* or *M. musculus* were aligned with reference sets with these settings for parameters: a length fraction of 0.35, a similarity fraction of 0.9, and mismatch, insertion, and deletion costs of 3 for each. Expression values were derived from uniquely mapped reads and normalized to total reads across all samples, without correction for reference sequence length. For comparisons across conditions and across species unique reads were further normalized as a ratio to *Gapdh* CDS reads in the same sample. These were further transformed as the natural logarithm of the ratios (73).

### Statistics

Graphs were generated using GraphPad Prism version 10.2.0, Python 3.11 in Jupyter Notebook (Project Jupyter; https://jupyter.org), or SYSTAT v. 13. Statistical analyses were performed using *R* v. 4.4.3, GraphPad Prism, or SYSTAT v. 13. Antibody titers and viral copy numbers were analyzed by ordinary one-way analysis of variance (ANOVA) followed by Tukey’s multiple comparisons test. Two-tailed Student’s t tests were used for comparisons of transcription values for individual genes between infection conditions. Means are reported with 95% confidence intervals. Box-whisker density plots featured either means with 95% confidence intervals or medians with 25th and 75th quartiles.

## Data availability

Sequencing data, including fastq format files of Illumina reads and associated sample descriptions (BioSamples), have been deposited in the NCBI SRA and BioSample databases under BioProjects PRJNA1026327 and PRJNA1026365. Accession numbers for Biosample identifications, Illumina reads (SRR) for lungs and brains, and Gene Expression Omnibus (GEO) data for lung and brain results are given in Table 4. The GEO accession number for the Third-Party Analysis of RNA-seq reads for *M. musculus* lungs from (45) is GSE296742. The descriptions and links for reference and data sets deposited with the Dryad Digital repository (http://dryaddata.org) are listed in Table S7.

**Table 4.** Data availability for RNA-seq reads and Gene Expression Omnibus data for lungs and brains of *P. leucopus*.

| ID | sex | group | day | exp | lung_Biosample | lung_SRR | lung_GEO | brain_Biosample | brain_SRR | brain_GEO |
| --- | --- | --- | --- | --- | --- | --- | --- | --- | --- | --- |
| 25013 | F | V | 3 | 1 | SAMN37756213 | SRR26353253 | GSM8967291 | SAMN37737300 | SRR26335496 | GSM8967261 |
| 25018 | F | C | 6 | 1 | SAMN37756214 | SRR26353252 | GSM8967292 | SAMN37737301 | SRR26335497 | GSM8967262 |
| 25025 | F | V | 3 | 1 | SAMN37756215 | SRR26353239 | GSM8967293 | SAMN37737302 | SRR26335498 | GSM8967263 |
| 25117 | M | V | 3 | 1 | SAMN37756216 | SRR26353228 | GSM8967294 | SAMN37737313 | SRR26335501 | GSM8967264 |
| 25124 | M | C | 3 | 1 | SAMN37756217 | SRR26353219 | GSM8967295 | SAMN37737303 | SRR26335499 | GSM8967265 |
| 25128 | F | V | 6 | 1 | SAMN37756218 | SRR26353218 | GSM8967296 | SAMN37737304 | SRR26335488 | GSM8967266 |
| 25130 | M | V | 3 | 1 | SAMN37756219 | SRR26353217 | GSM8967297 | SAMN37737305 | SRR26335489 | GSM8967267 |
| 25136 | M | V | 6 | 2 | SAMN37756228 | SRR26353244 | GSM8967305 | SAMN43066961 | SRR30252597 | GSM8967268 |
| 25152 | M | C | 3 | 2 | SAMN37756231 | SRR26353241 | GSM8967306 | SAMN43066964 | SRR30252580 | GSM8967269 |
| 25157 | M | V | 6 | 1 | SAMN37756220 | SRR26353216 | GSM8967298 | SAMN37737306 | SRR26335490 | GSM8967270 |
| 25158 | F | V | 6 | 1 | SAMN37756221 | SRR26353249 | GSM8967299 | SAMN37737307 | SRR26335491 | GSM8967271 |
| 25159 | M | C | 6 | 1 | SAMN37756222 | SRR26353248 | GSM8967300 | SAMN37737308 | SRR26335492 | GSM8967272 |
| 25160 | F | C | 3 | 1 | SAMN37756223 | SRR26353251 | GSM8967301 | SAMN37737309 | SRR26335493 | GSM8967273 |
| 25161 | M | V | 6 | 1 | SAMN37756224 | SRR26353250 | GSM8967302 | SAMN37737310 | SRR26335494 | GSM8967274 |
| 25163 | M | V | 6 | 2 | SAMN37756232 | SRR26353240 | GSM8967307 | SAMN43066965 | SRR30252579 | GSM8967275 |
| 25165 | M | V | 3 | 2 | SAMN37756233 | SRR26353238 | GSM8967308 | SAMN43066966 | SRR30252578 | GSM8967276 |
| 25167 | F | C | 6 | 1 | SAMN37756225 | SRR26353247 | GSM8967303 | SAMN37737311 | SRR26335495 | GSM8967277 |
| 25169 | M | C | 3 | 1 | SAMN37756226 | SRR26353246 | GSM8967304 | SAMN37737312 | SRR26335500 | GSM8967278 |
| 25219 | F | C | 6 | 2 | SAMN37756234 | SRR26353237 | GSM8967309 | SAMN43066967 | SRR30252577 | GSM8967279 |
| 25225 | F | C | 3 | 2 | SAMN37756235 | SRR26353236 | GSM8967310 | SAMN43066968 | SRR30252576 | GSM8967280 |
| 25226 | F | C | 3 | 2 | SAMN37756236 | SRR26353235 | GSM8967311 | SAMN43066969 | SRR30252575 | GSM8967281 |
| 25228 | F | V | 6 | 2 | SAMN37756237 | SRR26353234 | GSM8967312 | SAMN43066970 | SRR30252596 | GSM8967282 |
| 25255 | M | V | 3 | 2 | SAMN37756238 | SRR26353233 | GSM8967313 | SAMN43066971 | SRR30252595 | GSM8967283 |
| 25280 | M | C | 6 | 2 | SAMN37756240 | SRR26353231 | GSM8967314 | SAMN43066973 | SRR30252593 | GSM8967284 |
| 25281 | M | C | 3 | 2 | SAMN37756241 | SRR26353230 | GSM8967315 | SAMN43066974 | SRR30252592 | GSM8967285 |
| 25287 | M | C | 6 | 2 | SAMN37756243 | SRR26353227 | GSM8967316 | SAMN43066976 | SRR30252590 | GSM8967286 |
| 25298 | F | V | 3 | 2 | SAMN37756245 | SRR26353225 | GSM8967317 | SAMN43066978 | SRR30252588 | GSM8967287 |
| 25304 | F | V | 3 | 2 | SAMN37756246 | SRR26353224 | GSM8967318 | SAMN43066979 | SRR30252587 | GSM8967288 |
| 25306 | F | V | 6 | 2 | SAMN37756247 | SRR26353223 | GSM8967319 | SAMN43066980 | SRR30252585 | GSM8967289 |
| 25313 | F | C | 6 | 2 | SAMN37756248 | SRR26353222 | GSM8967320 | SAMN43066981 | SRR30252584 | GSM8967290 |

Abbreviations: F, female; M, male; exp, experiment; C, control; V, virus; SRR, Sequence Read Run, GEO, Gene Expression Omnibus

## Acknowledgments

We thank Lbachir BenMohamed, Jonathan Duong, Izabela Ibraim, Ilhem Messaoudi, and Allen Jankeel for their contributions of advice, assistance, or materials for the study.

The research was supported by an institutional grant from the University of California’s COVID-19 Basic, Translational and Clinical Research Funding Opportunity to DNF and AGB, NIH grants AI-136523 and AI-157513 to AGB, NIH grant AI-186970 to IC, and NIH grant NS-116835 to TEL.

This work also utilized resources of the UCI Genomics Research and Technology Hub, parts of which are supported by NIH grants to the Comprehensive Cancer Center (P30CA-062203) and the UCI Skin Biology Resource Based Center (P30AR075047), as well as to the GRT Hub for instrumentation (1S10OD010794-01and 1S10OD021718-01).

