## Supplemental Figures for "SARS-CoV-2 virus infection of *Peromyscus leucopus* demonstrates that infection tolerance is not limited to agents for which deermice are reservoirs"

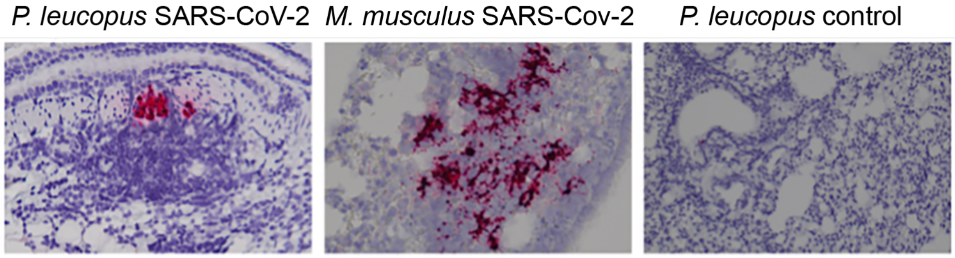


Figure S1. In situ localization of SARS-CoV-2 RNA in lung tissue by RNAscope hybridization. Formalin-fixed, paraffin-embedded lung sections from *Peromyscu leucopus* or *Mus musculus* and counter stained with hematoxylin were subjected to RNAscope in situ hybridization with a probe targeting SARS-CoV-2 genomic RNA. Red signal indicates the presence of viral RNA. The left panel is lung tissue of a *P. leucopus* animal inoculated with SARS-CoV-2 and at day 3 post-inoculation. It shows focal viral RNA localization within a peribronchial inflammatory infiltrate. The center panel is a lung section from K18-hACE2 transgenic *M. musculus* used as a positive control (reference 65), demonstrating widespread alveolar infection. The right panel is of lung tissue from uninfected *P. leucopus* used as a negative control and showing absence of hybridization signal.

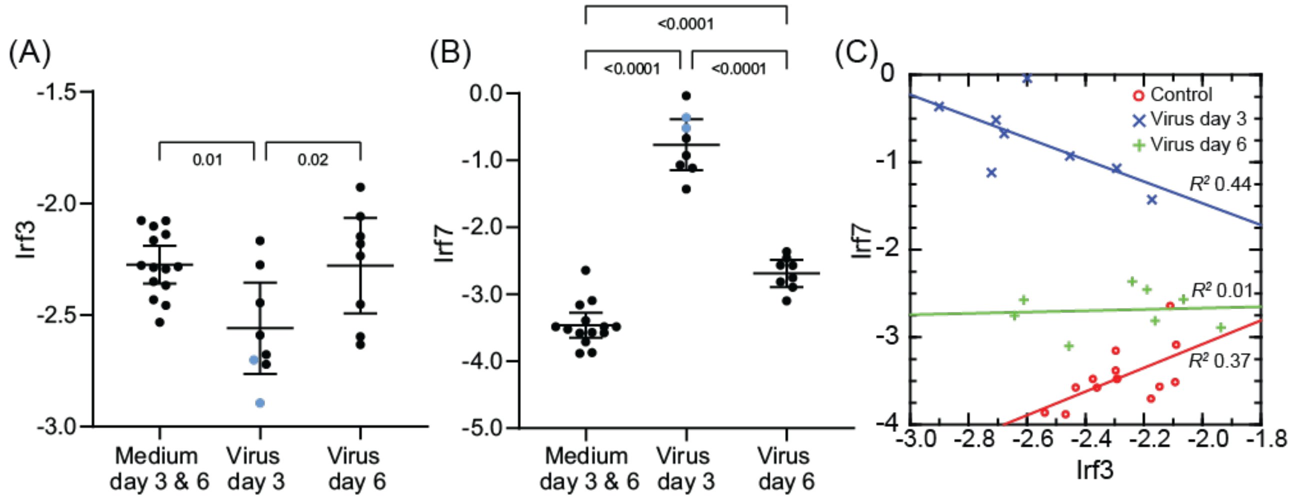


Figure S2. Transcription of the interferon regulatory factor genes *Irf3* and *Irf7* in the lungs of *P. leucopus* with (Virus) and without (Medium) infection by SARS-CoV-2 virus on day 3 or day 6 on infection. Panels A (*Irf3*) and B (*Irf7*) are box-whisker plots with means and 95% confidence intervals indicated. Panel C is a scatter plot with regression of transcripton of *Irf7* on *Irf3*. Transcription is shown as natural logarithm of ratiio of either *Irf3* or *Irf7* reads to *Gapdh* reads. Linear regressions with coefficients of determination ($R^2$) were determined separately for controls on days 3 and 6, virus-infected on day 3, and virus-infected on day 6. Data for graphs are provided in Table S3.

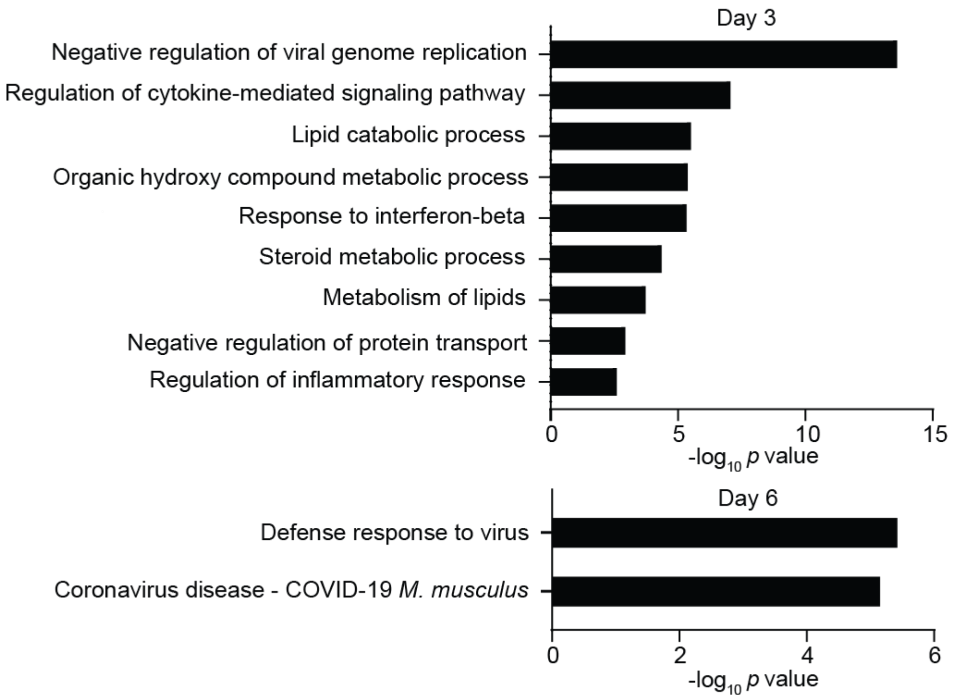


Figure S3. Functional enrichment analysis of differentially expressed genes of whole brains of *Peromyscus leucopus* infected with SARS-CoV-2 virus. Gene ontology (GO) analysis was based on upregulated genes with a false discovery $p$ value <0.05 and a $\log_2$ fold change of ≥1.5 for day 3 (upper panel) or day 6 dpi (lower panel). The data for the analysis are provided in Table S5. The identification numbers for each of the listed GO terms are given in Table S2.
